# Thermosensory neuron dysfunction alters transposable element regulation

**DOI:** 10.1101/2024.12.02.626368

**Authors:** Péter R. Szántó, David H. Meyer, Mélanie Bazin-Gélis, Thanh Vuong-Brender, Guy Teichman, Francy J. Pérez-Llanos, Daniel Rickert, Karl Köhrer, Oded Rechavi, Björn Schumacher

## Abstract

Transposable elements (TEs) can alter genome structure through transposition, and their activity is therefore tightly restricted by small RNA-mediated and chromatin-based silencing mechanisms. In *C. elegans*, elevated temperature can induce TE expression, raising the question of whether thermosensory neurons influence TE regulation. Here, we investigated the role of the AFD thermosensory neurons in TE regulation using multiple models of AFD dysfunction that altered TE expression; Mirage transposase induction emerged as the most reproducible phenotype across all AFD-dysfunctional strains. In the AFD triple mutant strain (PY9248), we observed strong Tc1 transposase expression, and genome-wide Tc1-enriched de novo insertions over generations. However, CRISPR reconstruction of the genotype in the N2 background did not reproduce the strong Tc1 phenotype, indicating that Tc1 activation and mutagenesis in PY9248 are background-associated rather than solely caused by loss of *gcy-8, gcy-18, gcy-23* function. These findings support a model in which AFD neuron dysfunction reproducibly alters TE expression, particularly of Mirage, while heritable Tc1-mediated mutagenesis requires an additional, currently uncharacterized factor present in the AFD triple mutant background.

## Introduction

Transposable element (TE) activity jeopardizes genome integrity as their insertions could lead to de novo mutations and structural changes in the genome. The potentially mutagenic TEs are therefore silenced by mechanisms such as regulatory small RNAs, particularly in the germline, to prevent harmful mutations that can occur if these elements insert into open reading frames or regulatory sequences^1,2^. In contrast to the genetic content, the epigenome of germ cells can be altered in response to environmental changes^3^. The ensuing transgenerational epigenetic changes are typically only maintained for a limited number of generations and do not alter the genetic information content itself ^3^.

Here, we employ the paradigm of temperature control of TE silencing in *C. elegans*, to investigate how environmental changes can affect the genetic information .^4–6^. The nematode has served as a pivotal model system to investigate the mechanisms of TE silencing^7–11^ and interactions between somatic tissues and the germline^12–14^. TE activity in germ cells is normally repressed by sRNA-mediated silencing^15–18^. At elevated temperatures, the Tc1 transposon has been shown to mobilize in germ cells^4,6^. Thus far, it has been unexplored whether this desilencing is a cell autonomous effect or a temperature sensing controlled mechanism. Environmental temperature status and change are sensed primarily by the pair of amphid sensory AFD neurons^19–22^ that control the animal’s heat shock response^23^. Under physiological conditions, the AFD neurons detect temperature via three receptor guanylyl cyclases, GCY-8, GCY-18, and GCY-23. Upon acute heat stress, these GCY enzymes are activated, producing cGMP that triggers downstream calcium-dependent cellular responses^24–26^. However, under prolonged exposure to elevated ambient temperatures (25 °C and above), AFD neurons adapt by reducing their intracellular calcium concentration instead of maintaining the GCY-cGMP-Ca² signaling axis activation ^19–21,27^. Here, we tested whether the status of AFD neurons might impact transposon silencing and thus regulate TE activity and mutagenesis.

We found that dysfunction of the AFD thermosensory neurons leads to TE desilencing and induced expression on transcriptomics level. In the original PY9248 AFD triple-mutant background, we established that activation of the Tc1 transposons leads to de novo genomic insertions and mutagenesis. However, subsequent validation experiments showed that this phenotype cannot be attributed solely to the triple mutation of the three AFD receptor guanylyl cyclases. Our data establish that the thermosensory AFD neurons regulate the repression of the Mirage transposons and raise new questions regarding the additional background factor that enables heritable Tc1 insertions through altered germline genome stability and mutagenesis.

## Results

### AFD neuron-dependent transposase expression induction

To assess TE expression, we first employed an RT-qPCR-based assay to measure transposase mRNA levels for selected TE families. We chose the Tc1 and Mirage TE families because they have been reported to be highly expressed in *C. elegans* mutant strains in which germline TE-silencing mechanisms are impaired^28^. To ensure accuracy, we targeted the perfect consensus region of each TE family with two independent primer pairs: Tc1(A), Tc1(B), Mirage(A), and Mirage(B) (**Supplementary Figure S1A**). To test the role of the AFD thermosensory neurons, we employed three distinct experimental systems of AFD impairment. First, we examined an AFD split-caspase ablation strain, hereafter referred to as AFD:SCA. This strain contains a transgenic construct expressing split caspases under the control of the *gcy-8* promoter, which induces apoptotic ablation of the AFD neuron.^22^. In AFD:SCA animals, Tc1 transposase expression mRNA mildly but not significantly increased, whereas Mirage transposase expression was significantly induced, exhibiting an approximately 9-fold increase in mRNA levels compared to wild-type (WT) controls **(Figure 1A, Supplementary Figure S1B)**. Raw data and statistical comparisons for each analysis are provided in the statistical source data supplementary file.

**Figure 1.**
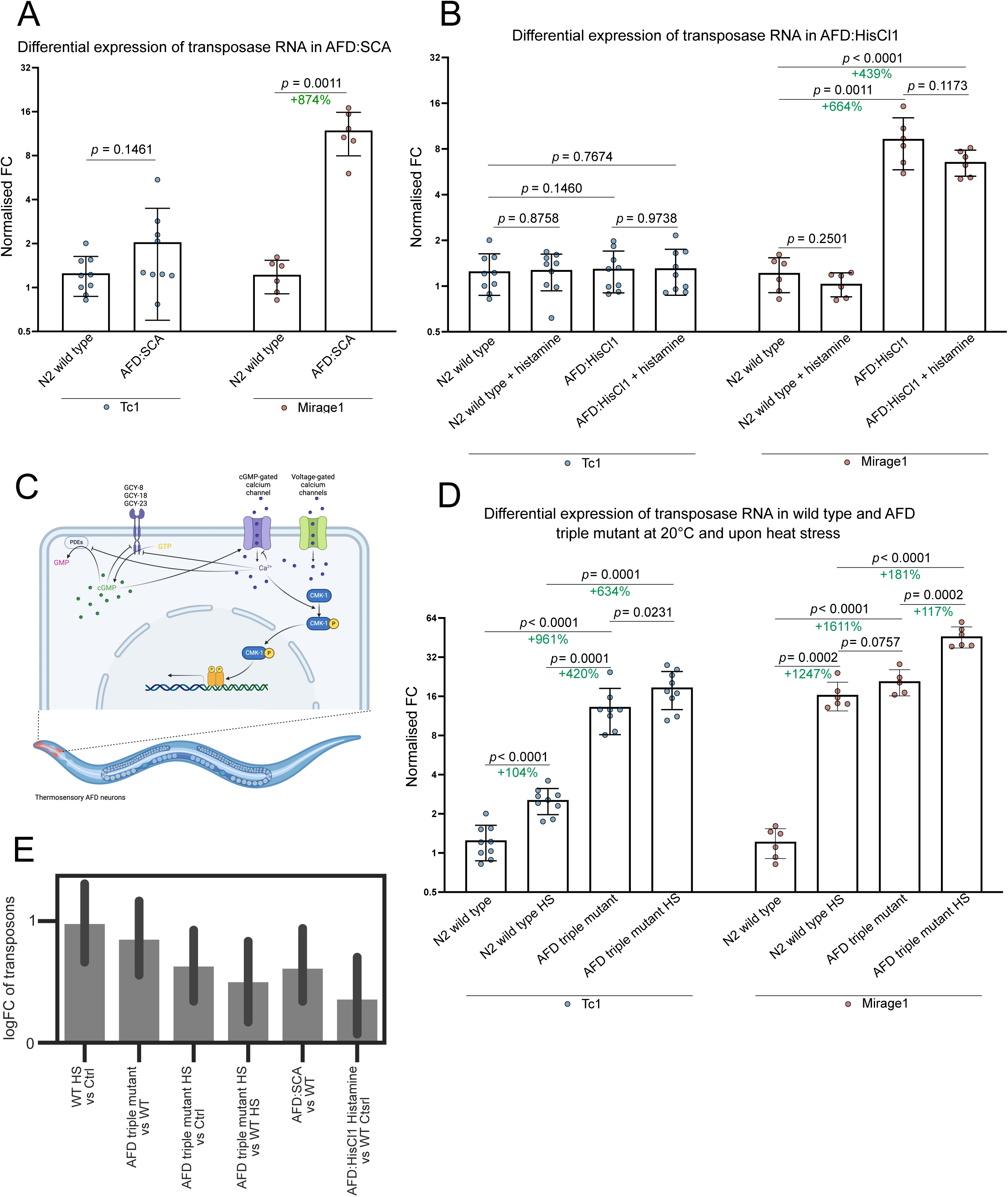
AFD dysfunction triggers transposon expression. RT-qPCR was performed to quantify the differential expression of transposase mRNA in various AFD neuron-depleted *C. elegans* strains. The experiment was repeated three times, each time with independently set up three independent cohorts consisting of 3000 nematodes. All biological replicate was measured in technical triplicates. Each data point represents the mean of a biological replicate, with data from these nine independent populations combined (ℕ=3, N=9). Target transposases, were measured with: Tc1 (B) for Tc1 transposase, and Mirage (B) for Mirage transposase. The Mirage (B) target was measure in two independent experiments, resulting in a total of six biological replicates. Relative quantities were calculated using the 2^ΔCt^ method and normalized using a normal factor derived from the relative quantities of the housekeeping genes tbg-1, act-1, eif-3.C, and rpl-32. This provided the normalized fold change (FC) values for the target transposases. Statistical significance was determined using an unpaired t-test with Welch’s correction, following normality assessment with the Shapiro-Wilk test. The p-values for the comparisons are indicated on the figure, along with the percentage change where relevant. A) Differential expression of transposase mRNAs in AFD split caspase-depleted strain (AFD:SCA) and wild type in comparison. B) Differential expression of transposase mRNAs in AFD histamine-gated chloride channel strain (AFD:HisCl1) compared to the wild type control. Populations were cultured on standard NGM and NGM supplemented with 1 mM histamine. C) Model of the thermosensory response pathway in the AFD neuron, adapted from ^26^. D) Differential expression of transposase mRNAs in AFD triple mutant strain compared to the wild type control, cultured under normal experimental temperatures and heat-stress conditions. E) The barplot shows the mean logFC of all transposable elements in WS291 in the respective pairwise comparisons. The error bar indicates the 95% confidence interval.

To test whether TE expression changes were reproducible using an independent method of AFD neuron manipulation, we utilized a transgenic strain expressing a histamine-gated chloride channel^29^ under the control of the *gcy-8* promoter, hereafter referred to as AFD:HisCl1. In this strain, the AFD neuron is hyperpolarized upon histamine treatment, rendering it functionally inactive. We performed RT-qPCR in AFD:HisCl1 animals under normal conditions and following supplementation with 1 mM histamine, which activates the chloride channel. In WT animals, histamine supplementation alone did not significantly alter Tc1 or Mirage transposase expression. In AFD:HisCl1 animals, Mirage transposase expression was significantly induced both under normal conditions and upon histamine supplementation, whereas Tc1 transposase expression remained unaffected (Figure 1B, Supplementary Figure S1C). The observed induction of Mirage transposase mRNA without histamine supplementation may be due to the previously reported leakiness of transgenic chloride channels, allowing Cl influx and causing neuron hyperpolarization even in the absence of histamine^29–31^.

To investigate more specifically the effect of functional impairment of AFD neurons on TE expression, we next examined animals carrying mutations in the three AFD-expressed receptor guanylyl cyclases, *gcy-8, gcy-18,* and *gcy-23*^24^. The strain used in these experiments was subsequently confirmed by genotyping to be PY9248, *gcy-8(oy44) gcy-18(nj38) gcy-23(oy150) IV*, rather than another triple mutant the also described in the literature, IK597, *gcy-8(oy44) gcy-18(nj38) gcy-23(nj37) IV*^24^. Therefore, unless stated otherwise, all references to the AFD triple mutant in this manuscript refer to the PY9248 strain. Both strains impair the same three AFD receptor guanylyl cyclase genes, although they differ in the *gcy-23* allele: *gcy-23(oy150)* introduces a premature stop codon^25,32^, whereas gcy-*23(nj37)* corresponds to a 954-bp deletion^24^. These three guanylyl cyclases are required for AFD-mediated thermosensory responses in *C. elegans*^24–26^. Reduced activity of these GCY enzymes is observed during adaptation after an initial temperature change, leading to reduced intracellular Ca² concentrations^19,20,26,33,34^. Loss of these GCY enzymes in the AFD triple mutant is expected to result in in a state of thermosensory perception and a neuronal state resembling chronic heat adaptation, despite maintenance at normal cultivation temperature ^25,33^. The AFD triple mutant animals displayed a similar induction of Mirage transposase as the two other AFD dysfunctional models above and in addition a marked induction of the Tc1 transposase (**Figure 1D, Supplementary Figure S1D**).

Given that various forms of heat stress have been reported to induce Tc1 activity in germ cells^6^, we further explored transposase regulation in the context of chronic heat stress. We compared Tc1 and Mirage transposase expression in young adult WT and AFD triple-mutant animals at the standard cultivation temperature of 20 °C and under chronic heat stress conditions, in which worms were cultivated at 25 °C and shifted to 26 °C at the L4 larval stage before collection in young adulthood (see Methods for details). We used 26 °C because this represents the lowest noxious level of chronic heat and is not yet toxic for AFD triple mutant animals, which develop normally at 25 °C and tolerate 26 °C from L4 onwards, whereas their development arrests when maintained at higher temperatures. Transposase mRNA levels from both TE families were significantly induced upon heat stress in WT nematodes, whereas AFD triple mutants showed strongly elevated Tc1 and Mirage expression at ambient temperature that was only slightly further elevated upon heat stress (**Figure 1D, Supplementary Figure S1D**).

To ensure that the observed TE desilencing was not confounded by the *mut-16(mg461)* background mutation, which has been reported in several *C. elegans* strains and is critical for TE silencing in germ cells^35^, we confirmed that the AFD triple mutant strain does not carry this mutation. (**Supplementary Figure S2**).

Because the three AFD perturbation models differ in how they impair AFD neuron function, we next performed an exploratory brood-size analysis to assess whether these strains also differed in reproductive fitness. The AFD triple mutant did not display a significant reduction in total egg number or viable progeny under standard conditions, although the number of unhatched eggs was increased **(Supplementary Figure S7A)**. In contrast, AFD:SCA animals showed a pronounced reduction in total egg number and viable progeny **(Supplementary Figure S7A)**. In the AFD:HisCl1 model, histamine treatment reduced total brood size and viable progeny, whereas histamine alone had no comparable effect in WT animals, supporting that adding histamine still has effect in activating the channel **(Supplementary Figure S7B)**. These results further support that the three AFD perturbation models differ not only in the mechanism of AFD impairment, but also in their broader physiological consequences.

To probe whether the AFD neurons could have a general effect on TE desilencing, we examined the transcriptome of the animals carrying dysfunctional AFD neurons. To comprehensively sequence transposon transcripts, we conducted total RNA sequencing (ribo-minus, see methods for details) in the three AFD neuron-impaired strains, i.e. AFD:SCA, AFD:HisCl1, and the AFD triple mutant, and, additionally, in the WT and AFD triple mutant under both normal temperature and chronic heat-stressed condition as applied above. Consistent with our RT-qPCR results, we observed a positive average induction across all TE families in all comparisons, including an additive effect of heat stress and the AFD triple mutant (**Figure 1E**). TEs showed a significant enrichment among the upregulated genes in the AFD:SCA strain, the AFD triple mutant, upon heat stress in WT worms, an additive upregulation upon heat stress in the AFD triple mutant, and a non-significant positive enrichment in the AFD:HisCl1 mutant upon Histamine treatment (**Supplementary Figure S3A-G**). Mirage showed the most consistent average upregulation and enrichment in all comparisons, while Tc1 showed the strongest effect in the AFD triple mutant and heat-stress, mirroring our RT-qPCR results (**Supplementary Figure S3H, I**).

Our results using three independent models of AFD dysregulation demonstrate that TE RNA expression is induced amid dysfunction of the thermosensory AFD neuron. However, the different types of AFD impairment distinctively affect the expression of the different TE families, likely due to their distinctive consequence on AFD function. The AFD:SCA model induces apoptosis and this strain exhibits severe impairments in fertility, development, and temperature resistance and shows lethality upon elevated temperature thus precluding heat stress experiments. The AFD:HisCl1 model induces hyperpolarization; however, the channel has been shown to be leaky, resulting in sustained hyperpolarization even in the absence of histamine^29,30^, complicating experimentation and requiring precise histamine control. In contrast, in the AFD triple mutant the neurons remain intact except for its thermosensory function ^19,20,33,34^. Furthermore, hierarchical clustering of TE family enrichment scores grouped the AFD triple mutant with heat-stressed wild-type transcriptomes, suggesting that the AFD triple mutant exhibits a TE expression profile similar to WT experiencing chronic heat stress, with further induction of TE expression upon additional heat stress. In contrast, AFD:SCA and AFD:HisCl1 formed a distinct outer group. (**Supplementary Figure S3I**). Together, the transcriptome analysis confirms that TE expression is induced both by AFD dysfunction and by heat stress. Consistently, the AFD triple mutant was reported to mount a systemic chronic stress response, as evidenced by increased peroxide resistance^32^. Based on these findings, we continued working with the AFD triple mutant as it appeared to model a neuronal state of chronic heat stress perception even at the normal cultivation temperature of 20 °C. We, therefore, proceeded to utilize the AFD triple mutant as our primary model for investigating the role of neuronal thermosensory regulation on TE activity.

### Disruption of TE regulation and the sRNA pathway in AFD triple mutant

To investigate the mechanistic basis of the TE induction in the AFD triple mutant, we analyzed the overall RNA sequencing data, which we further complemented with proteomics. First, we focused on gene expression changes that behave in a similar additive pattern as the average logFC of TEs (**Figure 1E**), i.e. genes that are significantly up-(respective down-) regulated upon heat-stress in WT, in the AFD triple mutant, and further upon heat-stress in the AFD triple mutant (**Figure 2A**). Intriguingly, *csr-1* and *ckk-1* were AFD mutant and heat-stress dependently repressed and induced, respectively, mirroring the TE expression levels in the AFD triple mutants and the additive effect of heat stress (**Figure 2A**). The Argonaute protein CSR-1 was shown to regulate gene silencing in the germ cells^36–38^, while the calmodulin kinase kinase CKK-1 operates in the thermosensory intracellular signal transduction in the AFD neurons. Consistent with the TE desilencing, a pathway enrichment analysis revealed significant enrichments in transgene expression and transposon silencing pathways in the additively downregulated genes (**Figure 2B**). The mechanisms responsible for silencing transgenes and transposons in the germline rely on a similar molecular machinery to recognize ’non-self’ transcripts^7,8,11,39^. The transgene silencing mechanisms that we find repressed here, were experimentally linked to the AFD neuron function in the accompanying manuscript by Teichman et al. Consistent with the role of sRNAs in silencing TEs, and foreign genetic elements in the germline, our transcriptome analysis revealed that numerous genes involved in sRNA biogenesis and processing are downregulated in the AFD triple mutant (**Figure 2C).**

**Figure 2.**
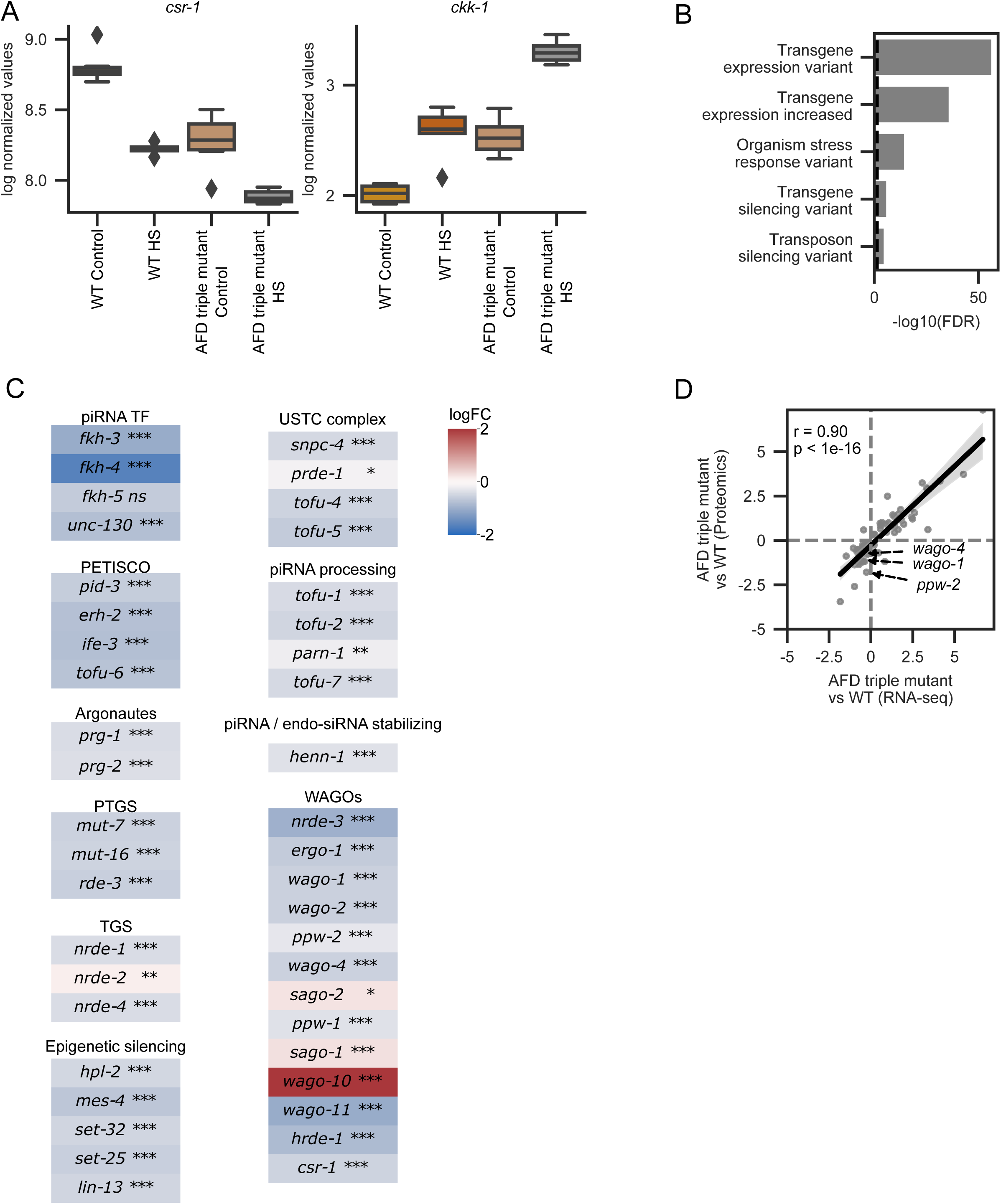
Transcriptome and Proteome analysis reveal desilencing mechanisms triggers in AFD triple mutants. Each strain and condition comprise five biological replicates, with independent nematode populations consisting of 3000 individuals (N=5). A) Two examples for additively changing genes upon heat stress (HS) in the wild type and the AFD triple mutant. The y-axis depicts the log normalized expression values. The boxplots show the quartiles of the datasets as boxes, the 1.5x interquartile range as the whiskers, and samples outside of this range as diamonds. B) Enriched pathways from an enrichment analysis with STRING^75^ for the additively downregulated genes, i.e. genes that are significantly downregulated in the AFD triple mutant, and upon HS irrespective of the strain. The x-axis shows the −log10 of the enrichment false discovery rate. For the full list see (**Supplementary Table 1**). C) Overview of genes involved in the regulation of sRNAs. The color code shows the logFC of the comparison AFD triple mutant vs. WT in the RNA-seq data. *<0.05, **<0.01, ***<0.001, ns>=0.05. D) Genes that are significantly regulated in the AFD triple mutant RNA-seq and proteomics are significantly positively correlated (Pearson correlation=0.9). The x-axis shows the logFC of the RNA-seq, the y-axis the logFC of the proteomics dataset.

In various organisms, TE silencing is mediated by the deposition of repressive histone modifications^40^. In line with the desilencing of TEs, the epigenetic silencing machinery (*hpl-2*, *mes-4*, *set-32*, *set-25*, and *lin-13*) is significantly downregulated in the AFD triple mutant (**Figure 2C**). To complement the transcriptomic analysis, we performed proteomic profiling of the AFD triple mutant and wild-type animals under both ambient and elevated temperatures, as described above. The transcriptomic and proteomic data showed a strong concordance, with a Pearson correlation coefficient of 0.9 indicating that the gene expression changes lead to similar responses on the proteome level (**Figure 2D**).

In conclusion, we found that many components of the sRNA machinery are downregulated in the AFD triple mutant at both the transcriptome and proteome levels. This is marked by the downregulation of sRNA biogenesis factors, sRNA processing genes, Argonaute proteins, and the post-transcriptional, as well as transcriptional gene silencing pathways, including the epigenetic silencing machinery. These data suggest that the AFD triple mutant is associated with reduced expression of multiple components of the sRNA and epigenetic machineries that silence TEs in the germline.

### Exploration of the pathway connecting the AFD neuron and germ cells

We next aimed to elucidate the signaling mechanism from the AFD neuron to the germline. First, we examined genes that have previously been reported to operate in or be influenced by the AFD neuron and to have an epistatic relationship with it ^41–43^. Consistent with compromised AFD function, we found those genes differentially expressed upon heat stress and in the AFD triple mutant (**Figure 3A**). Strikingly, we detected a pronounced induction of a range of neuropeptides in AFD triple mutants in line with a chronic thermosensory activation (**Figure 3B**). To further investigate the signaling mechanism through which the AFD neurons could impact on TE regulation, we computed a gene set enrichment analysis with publicly available differentially expressed gene sets with the significantly up-, respective down-, regulated genes in the AFD triple mutant (see Methods for details, **Supplementary Table 2**). The transcription factor *crh-1* was reported to be a major heat stress response regulator that mediates the expression of neuropeptides ^43^. In line with this reported role, the loss-of-function *crh-1(tz2)* mutant exhibits opposite patterns of gene expression compared to the AFD triple mutant (**Figure 3C, Supplementary Figure 4A**). A similar opposite gene expression pattern can be observed in loss-of-function *pmk-1(km25)* mutants (**Figure 3C, Supplementary 4 B**). PMK-1 is known to transmit somatic stress signals from the soma to germ cells via the secretion of SYSM-1 to regulate germ cell apoptosis and control stress-induced aneuploidy in the offspring^44^. Consistently, we observe a significant upregulation of the PMK-1 target gene *sysm-1* in the AFD triple mutant (logFC=1.31, FDR=7e-14).

**Figure 3.**
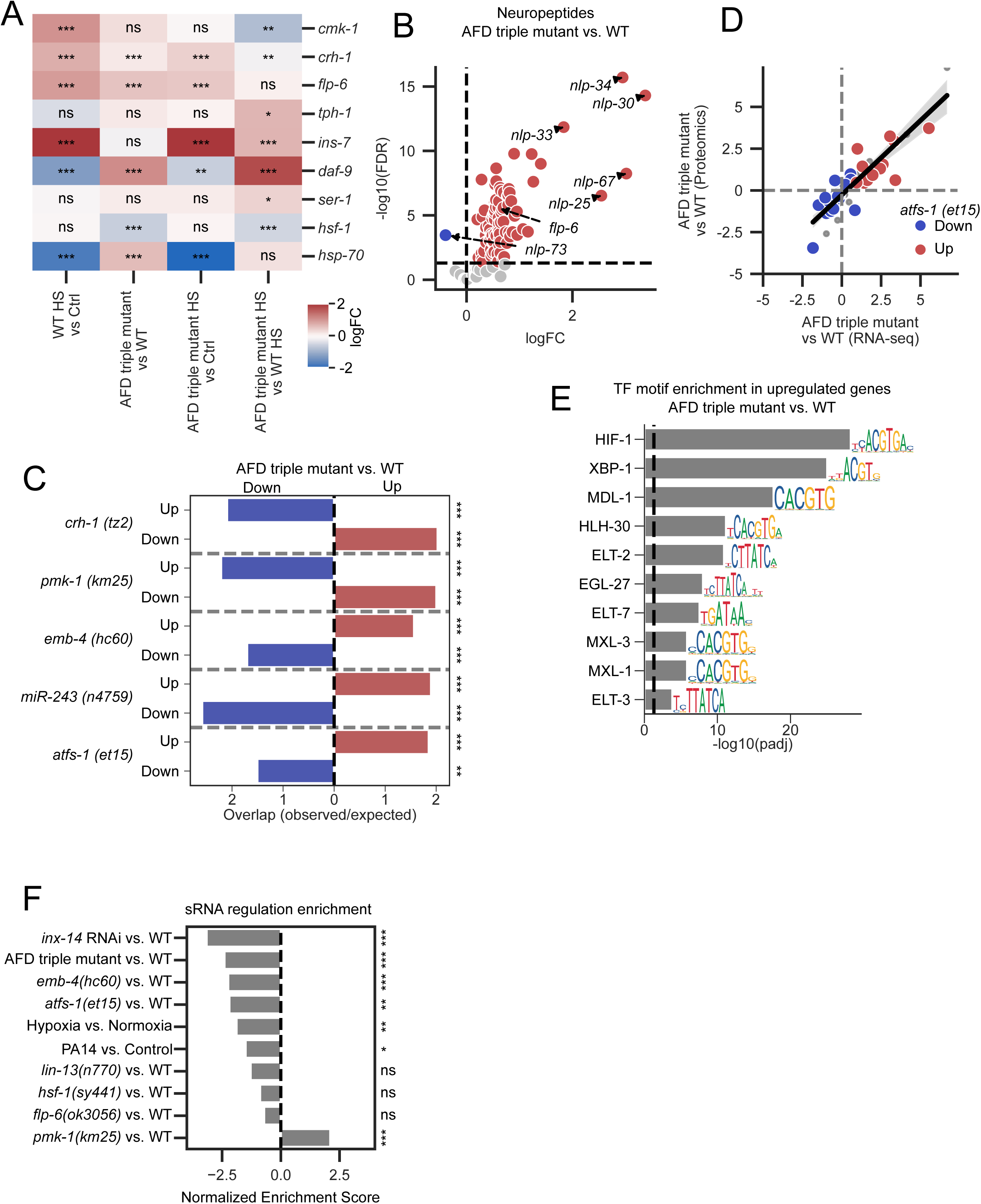
Transcriptome analysis identifies AFD-germline pathway regulating TE expression. A) Heatmap of logFC of the respective pairwise RNA-seq comparisons for genes known to be affected by the AFD neuron. The color code shows the logFC of the respective comparisons. *<0.05, **<0.01, ***<0.001, ns>=0.05 B) Neuropeptides are significantly upregulated in the AFD triple mutant compared to WT worms. The x-axis shows the logFC, the y-axis the −log10 false discovery rate. Red dots display significantly upregulated genes, blue dots significantly downregulated genes, and grey dots genes with an FDR >0.05. C) Comparison of publicly available differentially expressed gene sets with differentially expressed genes in the AFD triple mutant. WormExp^77^ was used to calculate the overlap of publicly available data with the significantly upregulated (red), respective downregulated (blue), genes in the AFD triple mutant. Five selected public datasets are shown, for the whole list see (**Supplementary Table 2**). The x-axis shows the overlap, i.e. the observed number of genes divided by the expected number of genes. *<0.05, **<0.01, ***<0.001 D) Genes regulated in the constitutive active atfs-1 (et15) mutant are similarly regulated in the AFD triple mutant. The correlation plot in Figure 3C is colored depending on whether the genes is significantly upregulated in the atfs-1(et15) mutant (red), significantly downregulated (blue), or not significantly changed (grey). E) The top 10 hits of a transcription factor motif enrichment analysis for the AFD triple mutant vs WT comparison (see method for details). The x-axis shows the −log10 of the multiple test adjusted p-value. The searched motif is depicted on the right side of each row. F) The results of gene set enrichment analyses for the sRNA regulation related genes from Figure 2D in public datasets. Each bar depicts the normalized enrichment score of the sRNA regulation related gene set in the specified dataset. A negative value indicates that the gene set is enriched in the downregulated genes. *<0.05, **<0.01, ***<0.001, ns>=0.05

In contrast, the loss-of-function *emb-4(hc60),* the loss-of-function *miR-243(n4759),* and the constitutive-active *atfs-1(et15)* mutants show a similar gene expression pattern as the AFD triple mutants (**Figure 3C, Supplementary Figure 4C-E**). The helicase *emb-4* has been reported to physically interact with the nuclear argonaute *hrde-1*, a critical component in the heritable silencing of genes and transposable elements. EMB-4/AQR facilitates this process by removing intronic barriers that otherwise impede the nuclear RNAi pathway, thereby enabling efficient silencing by HRDE-1 ^45^. This helicase has also been implicated in temperature tolerance in *C. elegans*^46^. miR-243 has been reported to interact with RDE-1, another argonaute, and plays a role in silencing endogenous RNA targets^47^. ATFS-1 is a transcription factor involved in the mitochondrial unfolded protein response and multiple stress response pathways, including the

*hif-1* mediated hypoxia response, *skn-1* mediated oxidative stress response, and *daf-16* mediated stress response^48^. Notably, genes regulated by ATFS-1 are enriched in the differentially expressed genes of the AFD triple mutant on the transcriptome as well as the proteome level (**Figure 3D, Supplementary Figure 4E**). Analyzing the transcription factor binding motifs present in the promoters of genes that are upregulated at the RNA level in our AFD triple mutant revealed a highly significant enrichment of HIF-1 transcription factor binding motifs, indicative of a role of HIF-1 in the mediator pathway (**Figure 3 E**).

To further investigate the potential mediators from the AFD neuron to the sRNA pathway in the germline, we analyzed relevant public datasets to assess the enrichment of sRNA pathway genes in relevant mutants and conditions (**Figure 3F, Supplementary Figure S5, S6**). In addition to the aforementioned mutants, we included a hypoxia treatment as a potent activator of HIF-1 ^49,50^; a treatment with the pathogen PA14 as an activator of PMK-1^51^; and a knock down of *inx-14*, which has been shown to mediate signal from the soma to the germline^52^. The majority of datasets exhibit significant downregulation of sRNA pathway genes, consistent with the downregulation observed in the AFD triple mutant, as shown in Figure 2E. Notably, only the *pmk-1(km25)* mutant dataset displays opposite sRNA pathway regulation behavior (**Figure 3F, Supplementary Figure S4, S5, S6**).

Based on our combined transcriptomic and proteomic analyses, comparisons to public datasets, and the genetic evidence for mediators of AFD-regulated germline transgene silencing in the recent manuscript by Teichman *et al.*^53^, we derived a plausible signaling model linking AFD neuron dysfunction to altered TE regulation in germ cells. In the AFD triple mutant, we observed an induction of *crh-1* and upregulation of neuropeptides. Downstream mediators include PMK-1^54,55^, a repressor of *hsf-1*, which is downregulated and functions as important regulator of the heat stress response in germ cells^56^. We also propose the involvement of *hif-1*, a key stress response mediator known to affect serotonin signaling ^57^, which has been reported to mediate the AFD neurogenic heat stress response to germ cells^42^, impacting *hsf-1* and chromatin maintenance^58^. Serotonin was shown to regulate HSF-1 activation in the germ cells in AFD neuron dependent manner^42^ and serotonin was also shown to activate *atfs-1*^59,60^, and constitutive active *atfs-1(et15)* mutant leads to downregulation of *emb-4*, which in turn was shown to mediate *hrde-1* dependent gene and transposon silencing specifically in germ cells^45^. These gene-regulatory changes are consistent with the genetic results of the recent manuscript by Teichman et al., which implicate AFD-mediated serotonin signaling in HSF-1-mediated regulation of the sRNA machinery in the germline.

### The AFD triple mutant displays inheritable Tc1-induced mutations

Since heat stress can lead to the de-silencing of TEs in germ cells^6^ and we detected a strong downregulation of the germline sRNA pathway (**Figure 2**), we hypothesized that the TE expression changes observed in the AFD triple mutant may affect the germline and potentially influence genome inheritance. As both our RT-qPCR (**Figure 1D**) and transcriptomic analyses (**Supplementary Figure S3H, I**) revealed an induction of Tc1 transposase RNA expression in the AFD triple mutant, we chose to investigate Tc1 insertion and its heritability. We utilized the well-established phenotype-based “twitching assay”, which has been validated in several studies as a reliable indicator of Tc1 transposon *de novo* insertion activity^61–63^. The twitching assay relies on spontaneous Tc1 transposon insertions into the *unc-22* locus, a 38 kb genomic sequence encoding the *C. elegans* homologue of the titin gene, essential for proper muscle function. When a Tc1 insertion disrupts UNC-22 function, nematodes display non-neurogenic muscle spasms that become evident upon nicotine treatment, which paralyzes WT animals whereas *unc-22::Tc1* insertion mutants continue to exhibit a twitching phenotype. ^61–63^. This assay serves as a robust genetic indicator of Tc1 transposon insertion activity, determined by the penetrance of the twitching phenotype in a given population derived from a single confirmed non-twitching founder hermaphrodite (**Figure 4A**).

**Figure 4.**
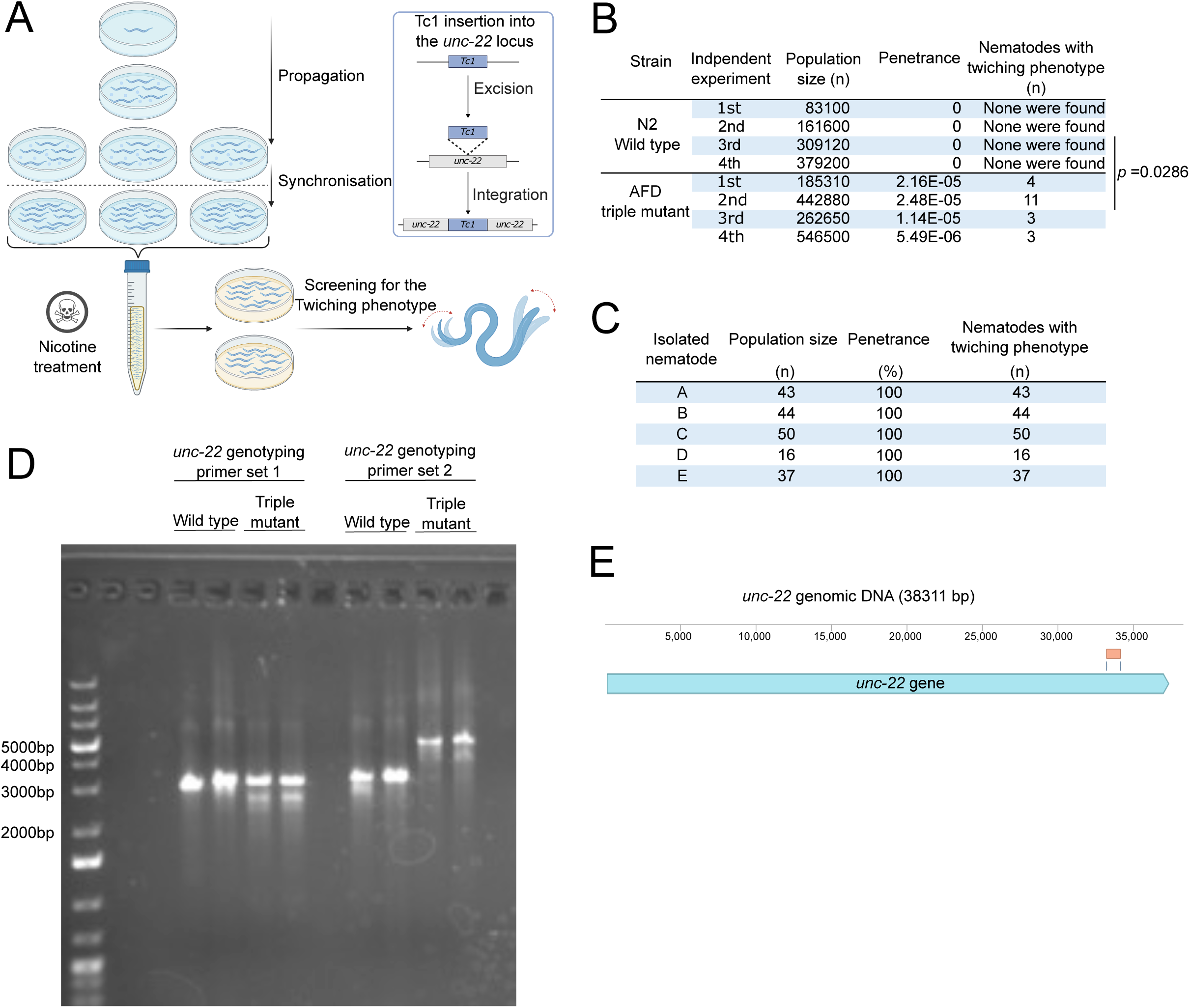
Phenotypic and genetic analysis shows *de novo* Tc1 insertion mutagenesis. A) Schematic representation of the experimental design and workflow for the twitching screen, including a description of the genetic background contributing to the observed twitching phenotype. B) Results of four independent experiments (=4) assessing the penetrance of the twitching phenotype in populations N2 wild-type and AFD triple mutant strains, derived from a single non-twitching mother. Data presented include population size (n), penetrance (expressed as ratio), and the number of nematodes displaying the twitching phenotype (n). The statistical comparison between the two strains was performed using the Mann-Whitney test, the p-value is indicated in the figure. C) Analysis of the inheritance pattern of the twitching phenotype in the F1 progeny from five individual worms (line A-E), isolated from the third and fourth independent experiments using the AFD triple mutant background. Penetrance of the twitching phenotype was assessed in these progenies. The F1 population size (n), penetrance (%) nematodes displaying the twitching phenotype (n) represented on the figure. D) Single-worm genotyping PCR analysis of the D-line isolate mother after the completion of egg-laying, as described in panel C. Seven primer pairs were used to genotype the *unc-22* genomic locus (38,311 bp). The figure shows the results for the first and second primer pairs, including wild-type control and AFD triple mutant D-line, with two technical replicates of the PCR. The primer set 2 revealed an insertion with expected wild-type product size of 3047 bp and a Tc1 transposon insertion size of 1032 bp. E) Schematic reconstruction of the *unc-22* genomic locus based on sequencing results. The annotation includes the *unc-22* gene (blue), Tc1 insertion (orange), and Tc1 flanking regions (dark blue lines), with a scale bar (bp) starting from the *unc-22* start codon.

While none of the WT animals displayed a twitching phenotype in each of the four independent experiments, we observed a penetrance of the twitching phenotype ranging from 5. 49 to 2.48 AFD triple mutant animals (**Figure 4B**). These results provide visually observable, phenotype-based evidence that elevated Tc1 transposase expression in the AFD triple mutant is associated with increased Tc1 transposon mobilization and mutagenesis. To assess whether the observed Tc1-associated mutations were stably inheritable, we independently recovered five twitching nematodes, established separate lines (A–E), and assessed the F1 progeny of these lines. The results demonstrated that the twitching phenotype was present in the F1 generation with 100% penetrance, confirming that the mutation is inheritable (**Figure 4C**).

To test whether the inheritable twitching phenotype was caused by a de novo Tc1 insertion, we performed single-worm PCR across the *unc-22* locus in one isolated line from the twitching screen, line D. We detected an insertion in the *unc-22* locus (**Figure 4D**). Sequencing of the PCR product confirmed the insertion of Tc1 into the coding region of the *unc-22* gene, 33,091 base pairs downstream of the start codon (**Figure 4E**). These findings indicate that the AFD triple mutant displays increased Tc1 transposon activity associated with a higher frequency of Tc1 insertion-induced inheritable mutations.

### The AFD triple mutant displays genome-wide Tc1-enriched de novo insertions

Following the observation of inheritable Tc1-associated mutations in the AFD triple mutant, we expanded our investigation to the genome-wide transposon-insertion landscape. We conducted a mutation accumulation experiment spanning thirty generations. We compared a founder P0 population with five independent F30 progeny lines, which were derived from the P0 population by transferring a single nematode before reproductive age in each generation, thereby minimizing the effects of natural selection. PacBio HiFi whole-genome sequencing (WGS) was used to analyze genomic changes and determine whether the AFD triple mutant displayed a higher frequency of mutation events compared to WT controls (**Figure 5A**).

**Figure 5.**
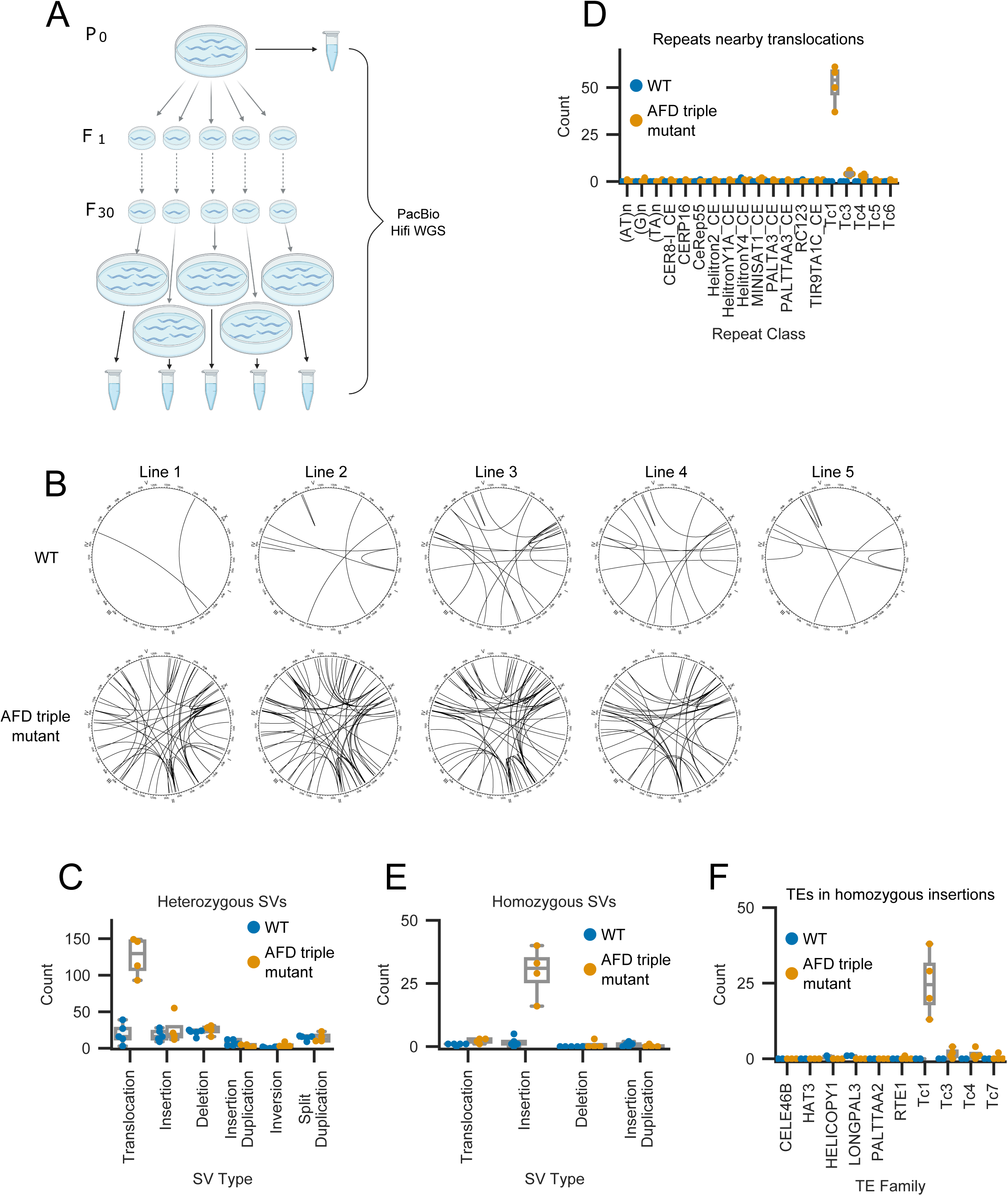
AFD neurons control inheritable TE insertion accumulation. A) Experimental design of the mutation accumulation experiment B) Circos plots showing *de novo* translocation events in the 5 independent F30 lines for WT (top row) and the AFD triple mutant (bottom row). Each line within the circles shows a translocation event that was not existent in the respective P0 sample. C) Quantification of heterozygous *de novo* structural variants (SV) in the WT F30 samples (blue), and the AFD triple mutant (orange). The y-axis shows the count of each SV type in the respective samples. Each dot is an independent repeat. D) Quantification of repeat element classes nearby (<10bp) the translocation break points. E) Quantification of homozygous *de novo* structural variants (SV) in the WT F30 samples (blue), and the AFD triple mutant (orange). The y-axis shows the count of each SV type in the respective samples. Each dot is an independent repeat. F) Quantification of transposable element classes in the homozygous inserted sequences.

To assess whether variants in known regulators of transposon activity could explain the observed phenotype, we first examined structural variants (SVs), small deletions (DELs), small insertions (INS), and single-nucleotide variants (SNVs) affecting genes known to influence transposon activity (**Supplementary Table 3, 4**). Although some genes in this list carried variants, these were primarily predicted to have modifier effects, were scattered across different strains, or were present in most samples, and therefore did not associate with the observed phenotype (**Supplementary Table 3, 4**). Having found no clear associating candidate variant among known TE-regulatory genes, we next analyzed the overall de novo mutation landscape after filtering out variants already present in the P0 generation of the respective strain (see methods for details).

First, we examined translocations and observed that the AFD triple mutant exhibited a marked increase in translocation number (**Figure 5B, C, E**). We confirmed the presence of the *gcy* mutations in the AFD triple mutant lines but had to omit one line that was WT for the *gcy* alleles and thus not a triple mutant. The heterozygous translocations in the AFD triple mutant were especially enriched to be nearby known Tc1 positions in the genome (**Figure 5D**). Further genomic analysis revealed that the AFD triple mutant strain also exhibited a strong increase in homozygous insertions compared to WT (**Figure 5E**). The inserted sequences were highly enriched in Tc1 transposons, indicating an increased Tc1 activity (**Figure 5F**).

Finally, we wondered why the *de novo* transposition events were specific to Tc1 but absent in other TE types despite their RNA expression. To assess whether the Tc1 transposon family might still be active in nematodes, we determined the number of transposon insertions in 152 wild isotypes of *C. elegans* from the Natural Diversity Resource (see methods for details). The *C. elegans* Tc1 transposon is related to the Mariner family of TEs, which is one of the most widely occurring TE type across species^64,65^. Consistently, the Tc1/Mariner family accounted for a large proportion of TE insertions in wild isolates, suggesting that this family has contributed substantially to recent TE accumulation in *C. elegans* genomes (**Supplementary Table 5**). In conclusion, our findings reveal pronounced Tc1-associated mutagenesis in the AFD triple-mutant, associated with Tc1 accumulation in heritable genomes over generations.

### CRISPR reconstruction of the PY9248 genotype separates Mirage induction from the PY9248-associated Tc1 phenotype

Given that the Tc1 activation phenotype was exclusive to the PY9248 AFD triple-mutant strain, in order to verify the genotype-phenotype causality, we sought to determine whether the observed TE expression phenotypes were reproducible in an independently generated strain carrying the same AFD receptor guanylyl cyclase mutations. To this end, we generated the BJS1456 strain, a CRISPR/Cas9-reconstructed AFD triple mutant in the N2 background carrying the same *gcy-8(oy44) gcy-18(nj38) gcy-23(oy150) IV* alleles as PY9248.

We then compared transposase expression in WT, the original PY9248 strain, our newly generated BJS1456, and the IK597 strain that carries the same *gcy-8* and *gcy-18* but a different *gcy-23(nj37)*alleles. We performed RT-qPCR for Tc1 and Mirage transposase expression under standard cultivation temperature and at 25°C and 26°C. The original PY9248 strain showed elevated Tc1 transposase expression at 20 °C and maintained higher Tc1 expression than the other tested strains across the temperature conditions, as before. In contrast, BJS1456 and IK597 did not reproduce the strong basal Tc1 expression observed in PY9248 and instead showed Tc1 expression profiles close to WT **(Figure 6A)**. These results indicate that the elevated Tc1 expression phenotype observed in the original PY9248 strain is not reproduced by CRISPR reconstruction and therefore cannot be attributed solely to impairment of the three receptor guanylyl cyclases.

**Figure 6.**
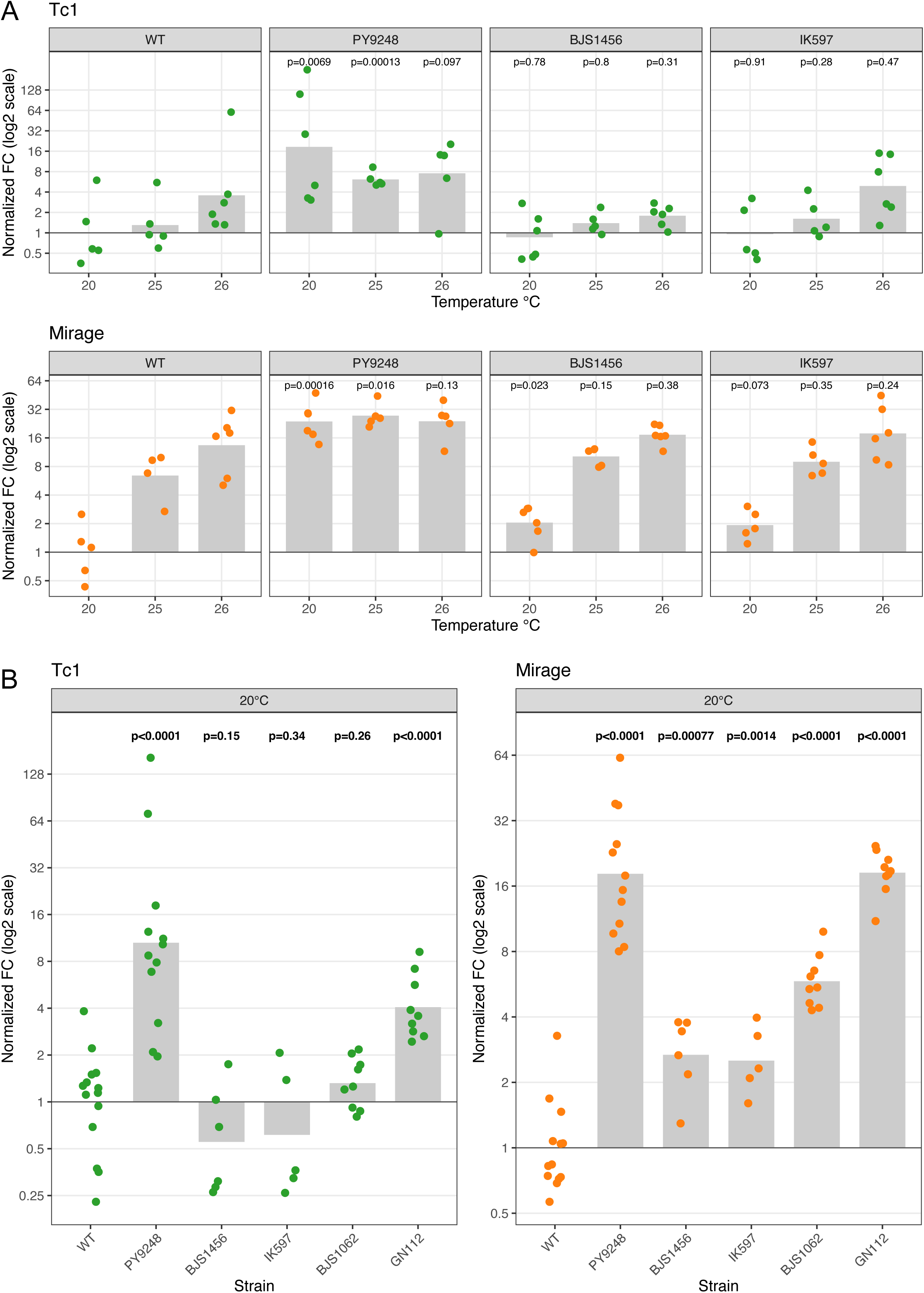
CRISPR reconstruction of the PY9248 AFD triple-mutant genotype separates Mirage induction from the PY9248-associated Tc1 phenotype. A) RT-qPCR of Tc1 (top) and Mirage (bottom) transposase expression in WT, original PY9248, CRISPR-reconstructed BJS1456, and IK597 animals at 20 °C or under chronic heat stress (25 °C, 26 °C) represented on x-axis. Expression is shown as normalized fold of change, relative to WT at 20 °C; gray bars indicate means and dots independent biological replicates. Each strain was compared with WT, within the given condition, by a paired t-test (p-value represented above each bar). B) Combined Tc1 and Mirage expression at 20 °C across AFD-perturbation models (WT, original PY9248, CRISPR-reconstructed BJS1456, IK597, BJS1062/AFD:HisCl1, and GN112/AFD:SCA). Each strain was compared with WT by Welch’s t-test (p-value represented above each bar).

Mirage transposase expression showed a different pattern. Although the original PY9248 strain displayed the strongest basal Mirage expression, Mirage induction was also detectable in the CRISPR-reconstructed BJS1456 strain and in the alternative AFD triple mutant IK597 **(Figure 6A)**. When all qPCR datasets were analyzed together, including the new Figure 6A data and the original Figure 1 data, the high Tc1 induction at 20 °C was specific to the PY9248 strain, however, AFD:SCA also showed statistically significant increase. In contrast, Mirage expression was significantly elevated across all AFD neuron dysfunction strains, reaching a level in AFD:SCA comparable to that in PY9248 **(Figure 6B)** which is consistent with the previous transcriptomics analysis of the strains **(Supplementary Figure S3I)**. These data separate the PY9248-associated Tc1 phenotype from the broader Mirage response observed upon AFD perturbation. Thus, the strong Tc1 expression and downstream Tc1-associated mutagenesis observed in the original PY9248 background should likely be interpreted as background-associated, whereas Mirage induction remains the reproducible TE expression phenotype linked to AFD neuron dysfunction.

## Discussion

TE expression is tightly controlled as TE mobilization can modify genomes through de novo insertions and structural variation^1,2^. Because heat stress can compromise TE silencing and promote Tc1 mobilization in *C. elegans* germ cells^4,6^, we investigated whether dysfunction of the AFD thermosensory neurons affects TE regulation.

Our initial results supported a model in which impaired AFD thermosensory signaling promotes TE desilencing, Tc1 activation, and heritable mutagenesis. In the PY9248 AFD triple-mutant strain used in our experiments, Tc1 and Mirage transposase expression were elevated, small RNA and epigenetic silencing pathways were transcriptionally repressed, and Tc1-associated phenotypic and genomic mutagenesis was detected. The twitching assay identified heritable *unc-22*-associated mutations, and genotyping PCR followed by sequencing confirmed a Tc1 insertion in the *unc-22* locus. Our mutation accumulation experiment and whole-genome sequencing further revealed increased structural variation and Tc1-enriched de novo insertions in the PY9248 background. These observations remain valid for the original PY9248 strain and demonstrate that this background displays elevated Tc1 expression and heritable Tc1-associated mutagenesis.

However, the validation experiments presented in Figure 6 refine this interpretation. CRISPR/Cas9 reconstruction of the PY9248 genotype in the N2 background, generating BJS1456 with the same *gcy-8(oy44), gcy-18(nj38), gcy-23(oy150)IV* alleles, did not reproduce the strong basal Tc1 expression observed in PY9248. Similarly, IK597, the other AFD triple mutant carrying *gcy-23(nj37)*, did not show the PY9248-like Tc1 phenotype. Thus, strong Tc1 expression and downstream Tc1-associated mutagenesis cannot be attributed solely to loss of the three AFD receptor guanylyl cyclases. The combined RT-qPCR analysis supports this conclusion: Tc1 induction was strongest in PY9248, while BJS1456 and IK597 remained close to WT. AFD:SCA showed at most a modest Tc1 increase, but clear heritable Tc1-induced mutagenesis and genome-wide Tc1 accumulation were detected only in PY9248. Therefore, AFD dysfunction may create a permissive state for TE transcriptional activation, while robust Tc1 activation and mobilization appear to require an additional factor present in the PY9248 background.

We examined our whole-genome sequencing, transcriptomic, and proteomic data to identify a candidate explanation for the PY9248-specific Tc1 phenotype. We did not identify variants in known TE regulators, small RNA pathway genes, or established Tc1-control factors that clearly segregated with the phenotype. These analyses do not exclude a causative background feature, since the relevant factor could be a noncoding variant, a structural variant with regulatory consequences, a cryptic modifier, altered Tc1 copy number, or an uncharacterized gene affecting Tc1 transcription, processing, or mobilization. Indeed, PY9248 contains a higher Tc1 copy number than the other strains tested, which may contribute to its elevated Tc1 activity. The most parsimonious interpretation is therefore that PY9248 carries an additional, currently unidentified permissive factor that enables strong Tc1 expression and mobilization, either independently or in interaction with the AFD triple-mutant physiological state.

The different outcomes of the AFD mutations in the distinct strain backgrounds narrow, but do not eliminate, the link between AFD neuron dysfunction and TE regulation. Across AFD dysfunction models, multiple TE families were affected, indicating that AFD perturbation and heat stress alter the broader TE expression landscape rather than acting exclusively on Tc1. Among these TE families, Mirage induction remained the most reproducible response. Mirage transposase expression was induced in all AFD neuron dysfunctional models (AFD:SCA, AFD:HisCl1, PY9248, BJS1456, and IK597). Thus, unlike Tc1, Mirage induction is not restricted to the PY9248 background and represents a robust TE-expression phenotype associated all the examined AFD neuron dysfunctional strains.

The biological meaning of Mirage induction remains unresolved. Because the RT-qPCR and transcriptomic analyses were performed on bulk whole-worm extracts, we couldn’t determine whether Mirage expression occurs in germ cells, somatic tissues, or both. If Mirage is induced in the germline, it may reflect altered germline TE silencing downstream of AFD dysfunction. If it is primarily somatic, it may instead represent a systemic stress-response output or tissue-specific TE transcriptional response. Moreover, TE expression does not necessarily imply TE mobilization. Even if Mirage is expressed in the germline, the transcripts may be nonfunctional, may lack the capacity to generate active transposase, or may be constrained by additional post-transcriptional regulatory mechanisms. Therefore, future work should dissect the tissue specificity and functional consequences of Mirage induction using germline-enriched RNA profiling, single-molecule RNA FISH, tissue-specific reporters, and gonad-specific qPCR.

The fertility phenotypes observed across AFD dysfunction models further suggest that AFD neuron perturbation can influence reproductive physiology, but the relationship between TE expression and fertility is not linear. PY9248 displayed the strongest TE induction and clear Tc1-associated mutagenesis, yet did not show a significant reduction in total brood size or viable progeny under standard conditions. In contrast, AFD:SCA and histamine-treated AFD:HisCl1 animals showed stronger fertility defects despite weaker or absent Tc1 activation. These observations argue against a simple model in which bulk TE expression alone explains reproductive impairment and suggest that fertility defects in these strains may also arise through TE-independent consequences of AFD neuron ablation or hyperpolarization.

In conclusion, our findings support a model in which AFD neuron dysfunction reproducibly alters TE expression, with Mirage induction representing the most robust and generalizable TE response across AFD neuron perturbation models. The original PY9248 AFD triple-mutant background additionally displays strong Tc1 transposase expression, heritable Tc1-induced *unc-22* mutations, and genome-wide Tc1-enriched de novo insertions over generations. However, because CRISPR/Cas9 reconstruction of the PY9248 genotype does not reproduce the strong Tc1 phenotype, the Tc1-associated mutagenesis observed in PY9248 should be interpreted as background-associated rather than as a direct consequence of *gcy-8, gcy-18, gcy-23* loss of function alone. We therefore conclude that AFD thermosensory dysfunction is linked to changes in TE regulation, but heritable Tc1-mediated mutagenesis requires an additional factor present in the PY9248 background. Identifying this factor may reveal a previously uncharacterized mechanism of germline TE regulation. In parallel, further dissection of AFD neuron-induced TE expression, particularly of the Mirage family, will clarify how thermosensory neuronal activity affects TE expression and whether this response reflects somatic stress signaling, germline-specific TE regulation, or both.

## Acknowledgements

We thank the Regional Computing Center of the University of Cologne (RRZK) for providing computing time on the DFG-funded High Performance Computing (HPC) system CHEOPS as well as support and the Zentrum für Informations-und Medientechnologie, especially the HPC team at Heinrich Heine University. Worm strains were provided by the National Bioresource Project (supported by The Ministry of Education, Culture, Sports, Science and Technology, Japan), the Caenorhabditis Genetics Center (funded by the NIH National Center for Research Resources, US), and the C. elegans Gene Knockout Project at the Oklahoma Medical Research Foundation (part of the International C. elegans Gene Knockout Consortium). We thank Wen Chen for providing the list with all transposable elements and their clustering into families for the WS291 reference.

O.R. and B.S. acknowledge funding from the Deutsche Forschungsgemeinschaft (DFG) project 437407415 (SCHU 2494/10-1) and the John Templeton Foundation Grant (61734), B.S. from the European Research Council (ERC-2023-SyG, 101118919), the Hevolution Foundation (HF-GRO-23-1199212-35), the Deutsche Forschungsgemeinschaft (Reinhart Koselleck-Project 524088035, FOR 5504 project 496650118, FOR 5762 project 531902955, SFB 1678, SFB 1607, CECAD EXC 2030 – 390661388, DFG-ISF project 561031107, ANR-DFG project 545378328, and the DFG project grants 558166204, 540136447, 496914708, 437825591, 437407415, 418036758), the José Carreras Leukemia Foundation, DJCLS 04 R/2023, the Deutsche Krebshilfe (70114555), and the Bundesministerium für Forschung, Technologie und Raumfahrt (BMFTR) X-AGE project 02NUK105B. This work was supported by the DFG Research Infrastructure West German Genome Center, project 407493903, as part of the Next Generation Sequencing (NGS) Competence Network, project 423957469. NGS and analyses were carried out at the West German Genome Center Düsseldorf. B.S. received the DFG grant SCHU 2494/7-1 as part of the DFG Sequencing call #1.

## Author contributions

*Conceptualization*: P.R.S., D.H.M., G.T., O.R., B.S.; *Experimentation*: P.R.S., M.B.G.; *Bioinformatics*: D.H.M., P.R.S.; *Data analysis*: P.R.S., D.H.M.; *PacBio sequencing experiment lead:* K.K.,F.J.P.-L; *PacBio data curation and bioinformatics analysis:* F.J.P.-L., D.R.; *Manuscript writing*: P.R.S., D.H.M, B.S.; *Visualization*: P.R.S., D.H.M; *Supervision*: B.S.; *Funding acquisition*: O.R., B.S.

## Materials and methods

### *C. elegans* culture and maintenance

Nematodes were cultured on nematode growth medium (NGM) agar plates. Strains were maintained at 15°C for medium-term storage, and long-term storage by freezing stocks at −80°C in freezing buffer. Prior to experiments, nematodes were acclimated to 20°C for at least two generations. All experiments were conducted at 20°C unless otherwise specified. In all cases during heat stress experiments, synchronized populations were cultured at 25 °C and at the L4 larval stage shifted to 26 °C until collection on the first day of adulthood. Developmental synchronization was achieved using a bleaching solution. Collection and washing of nematodes was done with M9 buffer. The protocols, including reagent and material compositions, were based on standard methods described in WormBook^66^.

### Generation of the BJS1062 “AFD:HisCl1”

To generate the BJS1062 “AFD:HisCl1” strain, a starved population of the BFF96 strain was subjected to gamma irradiation at a dose of 90 Gy, with a Biobeam 8000 containing a Cs137 radionucleotide source. This strain contains an extrachromosomal transgenic construct encoding a histamine-gated chloride channel, along with a transcriptional fusion mCherry reporter and a rol-6 selection marker, all driven by the AFD neuron-specific gcy-8 promoter *[pgcy-8::HisCl1::SL2::mCherry::rab-3 3’UTR, pgcy-8::GCaMP6::rab-3 3’UTR; rol-6(GOF)]*. Following irradiation, the progeny was screened over six generations to select worms displaying 100% identifiable mCherry fluorescence, indicating successful integration of the transgenic construct. The resulting strain was then backcrossed four times with the wild-type strain to ensure genetic consistency and remove potential background mutations, then confirmed again for the 100% penetrance of the mCherry fluorescence. As a result we established the BJS1062 strain, that carries the integrated *[pgcy-8::HisCl1::SL2::mCherry::rab-3 3’UTR, pgcy-8::GCaMP6::rab-3 3’UTR]* transgene.

### Generation of the BJS1456 strain

Sequential CRISPR genome editing was used to generate BJS1456 having mutations identical to PY9248, *gcy-8(oy44) gcy-18(nj38) gcy-23(oy150).* A protocol similar to the one described in (Ghanta., K.S and Mello., C.C, Genetics. 2020) was used. Briefly, a 20 µl CRISPR injection mix contains 0.5 µl Cas9 protein 10 µg/µl (IDT), guide RNA (2.8 µl crRNA 0.4 µg/µl and 5 µl tracrRNA 0.4 µg/µl, IDT), 2.2 µl of a single strand oligonucleotide 1 µg/µl (IDT) or 500 ng of melted dsDNA as repair template and 800 ng co-injection marker plasmid pRF4. To generate *gcy-8(oy44),* a crRNA with the variable region AAGGATTGTGCATGACATAT was used; the repair template was made by amplifying the gDNA of PY9248 with forward GGCCATCAGTTGATAGGAAAATGC and reverse GAGAACATTGGATTCACGGACCA primers. To create *gcy-18(nj38),*a crRNA with the variable region AAGTTGAGCAATATACAGAC was used; the repair template was made by amplifying the gDNA of PY9248 with forward CGCACTGAAAGCCGTACGAAT and reverse ACCGTAATCCGACAGCTTAACCA primers. To generate *gcy-23(oy150),* a crRNA with the variable region GACTACACCACATTGATTAT and a single-strand DNA with the sequence ATGAGCCAGCATGCGGCTTCCGAAATGAAAAATGCGACTACATTACATAAATAAATAATGATTATCGGAGGATGCATTGTACTTCTGATCATTCTCCTTA were used. The injection mix was centrifuged for 5 min at maximum speed in an Eppendorf table-top centrifuge, then 0.5 µl of the mix was loaded into a glass needle. The glass needles were pulled using glass capillaries (World Precision Instruments, 1B100F-4) with a PC-100 Narishige Puller. The injection was performed using a Zeiss Axio Observer.A1 microscope with a Zeiss Neofluar 40X/0.75 objective and an Eppendorf micromanipulator InjectMan NI 2. Day one hermaphrodites were placed on a coverslip with dried 2% agar using a drop of halocarbon oil 700 (Sigma). The tip of the needles was broken using a manually-pulled glass fiber. 20 to 30 P_0_ hermaphrodites were injected for each CRISPR. F_1_ roller progenies were isolated and left overnight to lay eggs then genotyped on the following day using PCR and Sanger sequencing. The resulting strain was outcrossed twice.

### RNA Extraction for RT-qPCR and RNA sequencing

All RNA extractions for this study were performed using the following method. Worms were collected in a tube and washed with M9 buffer, followed by the addition of 1 mL Trizol® reagent.

The sample was homogenized by freeze cracking, which involved dropping it into liquid nitrogen and then melting it in an incubator. Subsequently, 100 μL of bromochloropropane (BCP) was added for phase separation. The sample was incubated at room temperature for 2 minutes and then centrifuged at 12,000 rcf for 15 minutes at 4°C. The upper, clear phase (∼600 μL) was collected into a fresh RNase-free tube, mixed with an equal volume (∼600 μL) of 70% ethanol, and vortexed.

The mixture was then processed using the Qiagen RNeasy Mini Kit® according to the manufacturer’s instructions, including on-column DNase digestion. The RNA was eluted in 50 μL of RNase-free water. The extracted RNA was quantified and its integrity assessed using Thermo Scientific Nanodrop spectrophotometer to ensure suitability for downstream applications.

### cDNA Synthesis

The SuperScript® III Reverse Transcriptase (Invitrogen) kit was used for reverse transcription reactions of isolated RNA samples. cDNA synthesis was performed in a 20 µL reaction mixture. The initial reaction mixture included 12 µL of RNA solution (1 µg RNA), 1 µL of random hexamer, and 1 µL of dNTP. This mixture was incubated at 65°C for 5 minutes and then subjected to a brief centrifugation. Following this, the mixture was placed on ice for 5 minutes.

Subsequently, 4 µL of 5x First Strand Reaction Buffer, 1 µL of DTT, 1 µL of RNase Out Recombinant RNase Inhibitor, and 1 µL of SuperScript III RT (200 U/µL) were added to the mixture. The contents were gently mixed by pipetting. The reaction was then incubated under the following conditions: 25°C for 5 minutes, 50°C for 60 minutes, and 70°C for 5 minutes.

Following the incubation, the cDNA was diluted to a final volume of 100 µL by adding 80 µL of double-distilled water (ddH2O). The resultant cDNA was evaluated using a control PCR to verify successful synthesis.

### Quantitative Real-Time PCR (qPCR)

Quantitative Real-Time PCR was performed using the PowerUp™ SYBR™ Green Master Mix (Thermo Fisher Scientific). The reaction mixture consisted of 10 μL of 2X PowerUp™ SYBR™ Green Master Mix, 9 μL of 1.1 μM forward and reverse primers, and 0.25 μL of cDNA template, with the total reaction volume adjusted to 20 μL using nuclease-free water. Each sample was measured in biological triplicates from each technical replicate.

A Template Master Mix was prepared by combining 10 μL of PowerUp™ SYBR™ Green Master Mix, 0.25 μL of cDNA template (50 ng/μL), and 0.1 μL of nuclease-free water per reaction, yielding a total volume of 11 μL per well. Separately, a Primer Master Mix was prepared at a concentration of 1.1 μM by mixing 5.5 μL each of 100 μM forward and reverse primer stocks with 489 μL of nuclease-free water to reach a final volume of 500 μL.

Each well of the qPCR plate was aliquoted with 11 μL of Template Master Mix and 9 μL of Primer Master Mix. The qPCR assays were conducted using a CFX Connect and CFX96 Touch real-time PCR detection system (BioRad). The expression levels of target genes were normalized to the reference genes *tbg-1, act-1, eif-3.C*, and *rpl-32*. Data acquisition and analysis were performed using CFX Manager Software (BioRad), with subsequent data processing carried out in Microsoft Excel 2019, R version 4.3.2, and GraphPad Prism 9.

### Total RNA sequencing

Libraries were prepared from 500ng total RNA. Enzymatic depletion of ribosomal RNA with the Illumina Ribo-Zero Gold rRNA Depletion Kit was followed by library preparation with the Illumina TruSeq Stranded Total RNA sample preparation kit. The depleted RNA was fragmented and reverse transcribed with random hexamer primers, second strand synthesis with dUTPs was followed by A-tailing, adapter ligation and library amplification (15 cycles). Next library validation and quantification (Agilent Tape Station) was performed, followed by pooling of equimolar amounts of library. The pool itself was then quantified using the Peqlab KAPA Library Quantification Kit and the Applied Biosystems 7900HT Sequence Detection System and sequenced on an Illumina NovaSeq6000 sequencing instrument with an PE100 protocol aiming for 30 million clusters per sample.

### Proteomics sample preparation

Whole-animal protein extracts were prepared from a homogenized population of 3,000 C. elegans. Samples were lysed in 8 M urea buffer with 50 mM triethylammonium bicarbonate (TEAB), supplemented with a 50x protease inhibitor cocktail (Roche). For chromatin digestion, Benzonase HC nuclease was added at a concentration of 25 units, followed by incubation at 37°C for 30 minutes to ensure complete chromatin digestion. The lysates were centrifuged at 20,000 × g for 15 minutes to remove cell debris, and protein concentrations were measured using a standard assay. For reduction and alkylation, 50 µg of protein was transferred to a 1.5 mL microcentrifuge tube. Dithiothreitol (DTT) was added to a final concentration of 5 mM, followed by incubation at 25°C for 1 hour. Chloroacetamide (CAA) was then added to a final concentration of 40 mM, and the samples were incubated in the dark for 30 minutes. Trypsin was then added at a 1:75 enzyme-to-substrate ratio, and the samples were incubated overnight at 25°C for complete proteolysis. On the following day, the enzymatic digestion was stopped by acidification with formic acid to a final concentration of 1%. The samples were subsequently purified using SDB-RP StageTips packed with two layers of SDB-RPS discs in 200 µL pipette tips. The StageTips were equilibrated by sequential centrifugation at 800 × g for 1 minute with 20 µL methanol, 20 µL buffer B (0.1% formic acid in 80% acetonitrile), and twice with 20 µL buffer A (0.1% formic acid in water). Acidified samples were centrifuged, and the supernatant was loaded onto the StageTips, followed by washing with buffer A and buffer B. The StageTips were then dried and stored at 4°C and submitted to the CECAD Proteomics Facility. All reagents mentioned in this preparation were provide by the CECAD Proteomics Facility (https://proteomics.cecad-labs.uni-koeln.de/home).

### Proteomics Data Acquisition

Samples were analyzed by the CECAD Proteomics Facility on an Orbitrap Exploris 480 (Thermo Scientific, granted by the German Research Foundation under INST 1856/71-1 FUGG) mass spectrometer equipped with a FAIMSpro differential ion mobility device that was coupled to an UltiMate 3000 (Thermo Scientific). Samples were loaded onto a precolumn (Acclaim 5μm PepMap 300 μ Cartridge) for 2 min at 15 ul flow before reverse-flushed onto an in-house packed analytical column (30 cm length, 75 μm inner diameter, filled with 2.7 μm Poroshell EC120 C18, Agilent). Peptides were chromatographically separated at a constant flow rate of 300 nL/min and the following gradient: initial 6% B (0.1% formic acid in 80 % acetonitrile), up to 32% B in 72 min, up to 55% B within 7.0 min and up to 95% solvent B within 2.0 min, followed by column wash with 95% solvent B and reequilibration to initial condition. The FAIMS pro was operated at −50V compensation voltage and electrode temperatures of 99.5 °C for the inner and 85 °C for the outer electrode.

For the Gas-phase fractionated library, a pool generated from all samples was analyzed in six individual runs covering the range from 400 m/z to 1000 m/z in 100 m/z increments. For each run, MS1 was acquired at 60k resolution with a maximum injection time of 98 ms and an AGC target of 100%. MS2 spectra were acquired at 30k resolution with a maximum injection time of 60 ms. Spectra were acquired in staggered 4 m/z windows, resulting in nominal 2 m/z windows after deconvolution using ProteoWizard ^67^.

For the samples, MS1 scans were acquired from 399 m/z to 1001 m/z at 15k resolution. Maximum injection time was set to 22 ms and the AGC target to 100%. MS2 scans ranged from 400 m/z to 1000 m/z and were acquired at 15 k resolution with a maximum injection time of 22 ms and an AGC target of 100%. DIA scans covering the precursor range from 400 - 1000 m/z and were acquired in 60 x 10 m/z windows with an overlap of 1 m/z. All scans were stored as centroid.

### Proteomics Sample Processing

The gas-phase fractionated library was build using DIA-NN 1.8.1 (Demichev 2020) using C. elegans WormBase database (PRJNA13758, downloaded 24/01/23) with settings matching acquisition parameters.

Samples were analyzed in DIA-NN 1.8.1 as well using the previously generated library and identical database. DIA-NN was run with the additional command line prompts “—report-lib-info” Further output settings were: filtered at 0.01 FDR, N-terminal methionine excision enabled, maximum number of missed cleavages set to 1, min peptide length set to 7, max peptide length set to 30, min precursor m/z set to 400, max precursor m/z set to 1000, cysteine carbamidomethylation enabled as a fixed modification. Afterwards, DIA-NN output was further filtered on library q-value and global q-value <= 0.01 and at least two unique peptides per protein using R (4.1.3). Finally, LFQ values calculated using the DIA-NN R-package. Afterwards, analysis of results was performed in Perseus 1.6.15^68^.

### Mutation Accumulation Experiment Design and Strain Handling

For this experiment, a single mother was isolated from each of the strains. These isolated mothers were used to propagate populations, which were synchronized by bleaching. This synchronized population served as the P0 founder population for the experiment. The populations were collected and stored at −80°C.

From these initial populations, five mothers were isolated and placed on separate plates, serving as the founding mothers of the experiment (N=5). Throughout the experiment, one L4 stage larva from each strain was transferred individually to a new plate in every generation, spanning 30 generations.

Upon reaching the F30 generation, a population was propagated and synchronized via bleaching to collection for whole genome sequencing (n=3000). All the synchronized populations were collected at the young adult stage by washing with M9 buffer and stored at −80°C, and frozen nematode stocks were established for long-term storage.

### High Molecular Weight Genomic DNA Extraction from *C. elegans*

High molecular weight (HMW) genomic DNA was extracted from *C. elegans* using the QIAGEN Puregene Cell Kit. Approximately 3000 nematodes were grown on NGM agar plates with E. coli OP50 strain and harvested using M9 buffer. The harvested worms were stored at – 80°C until DNA extraction. The extraction process commenced by adding 3 mL of Cell Lysis Solution and 15 μL of Proteinase K to the frozen worm pellet, allowing it to thaw. Once thawed, the pellet was resuspended by pipetting with a 1 mL wide-bore tip. The sample was incubated at 50°C for 2 hours, with gentle inversion of the tube every 30 minutes to ensure homogeneity.

Following incubation, 15 μL of RNase A was added, and the mixture was incubated at 37°C for 30 minutes. Subsequently, 1 mL of Protein Precipitation Solution was added, and the tube was mixed by inversion 15-20 times before being placed on ice for 2 minutes. The sample was centrifuged at 2000 x g for 10 minutes, and DNA was precipitated from the supernatant using 3 mL of isopropanol. The tube was gently inverted 50 times and centrifuged at 2000 x g for 5 minutes. The supernatant was discarded, and the DNA pellet was washed with 3 mL of 70% ice-cold ethanol. The tube was gently inverted to mix and then centrifuged at 2000 x g for 2 minutes. The supernatant was discarded, and any remaining ethanol was removed using sterile paper wipes. The DNA was eluted in 600 μL of TE buffer and incubated overnight at room temperature to maximize yield. The quantity and quality of the extracted DNA were assessed using a Thermo Scientific Nanodrop spectrophotometer and the Agilent TapeStation System.

### PacBio whole genome sequencing

#### HiFi Whole Genome Library Preparation

DNA quality and quantity were assessed using a NanoDrop spectrophotometer, the Qubit BR DNA assay (ThermoFisher), and the Femto Pulse system (Agilent) with the FP-1002 protocol (165 kb) to evaluate fragment size distribution.

Long-read whole-genome sequencing libraries were prepared using the HiFi low-input protocol as described in the SMRTbell® Prep Kit 3.0 (Version 02; March 2023). Genomic DNA (1,500– 7,900 ng) in Low TE buffer was used as input for library preparation. DNA shearing was performed with a MegaRuptor3 (Diagenode) using long hydropores at a shearing speed of 27– 30, targeting a fragment size of 10–15 kb when appropriate. Samples with a pre-sheared average fragment size suitable for library preparation were not further processed. Sheared DNA was purified using the DNeasy PowerClean Pro Cleanup Kit (Qiagen) according to the manufacturer’s instructions, eluting the DNA in 50 µl of water. For samples with limited DNA, up to three sequential 50 µl elutions were performed on the same column, with eluates pooled to minimize DNA loss.

Pooled samples were concentrated to a final volume of 50 µl using a SpeedVac at room temperature. DNA quantification was performed at all steps using the Qubit dsDNA assay (ThermoFisher). Fragment size distribution was assessed using a Femto Pulse (Agilent) with the FP-1002 protocol (165 kb) by diluting the sheared DNA to 0.1 ng/µl. Sheared samples were further purified with 1X SMRTbell cleanup beads (PacBio), washed twice with 80% ethanol, and eluted in 47 µl of Low TE buffer (PacBio). DNA concentration was again determined using the Qubit dsDNA HS assay (Thermo Fisher). For library preparation, 46 µl (480–4,000 ng) of each sample was used, following the manufacturer’s protocol. This process included DNA repair, A-tailing, adapter ligation, and a nuclease treatment step. AMPure bead size selection was performed to remove fragments smaller than 5 kb, and final libraries were eluted in 11–15 µl of Elution Buffer (PacBio).

A 1:10 dilution of the eluted library was prepared for quality control. Library concentration was measured using Qubit (Thermo Fisher), and size distribution was analyzed with the Femto Pulse (Agilent) using the FP-1002 protocol. Four to six libraries were pooled equimolarly based on their average fragment size and total DNA amount, with each library containing a unique barcode. The final library pool concentration was verified using the Qubit dsDNA HS assay (Thermo Fisher), and fragment size distribution was confirmed on the Femto Pulse. A total of seven final library pools were prepared and stored at 4°C until sequencing.

#### HiFi Whole Genome Single Molecule Real-Time Sequencing

Sequencing of each SMRTbell library pool was conducted following PacBio’s standard sample preparation and sequencing guidelines (SMRT Link version 11.1.0.166339). Sequencing primers and polymerase were bound to each library pool using the Sequel® II Binding Kit 3.2 (PacBio). Polymerase-bound SMRTbell complexes were purified, achieving a recovery rate of ≥75%, as confirmed using the Qubit dsDNA HS assay (Thermo Fisher). Each library pool, including internal control, was sequenced on an SMRT Cell 8M with an on-plate loading concentration of 70–85 pM. Run conditions included a 120-minute pre-extension time and an 1800-minute movie time. Sequencing was performed on Sequel II and Sequel IIe systems.

#### PacBio Data Analysis

Raw subreads were obtained after filtering polymerase reads and then subjected to Circular Consensus Sequencing (CCS) analysis resulting in HiFi reads using SMRT Link (version 11.1.0.166339 and 13.0.0.207600). HiFi reads were further demultiplexed using Lima with a barcode score threshold of 96. On average, the sequencing generated ∼360,000 HiFi reads per genome with a quality value of ≥20, an N50 read length of approximately 8.6 kb, a predicted base accuracy of 99%, and a minimum of three passes per molecule. Datasets were organized into generations: P0, F30 for the wild type and AFD triple mutant respectively, with one P0 sample rep and five samples corresponding for the F30 descendants of the five independent lines. Each analysis was performed individually on sample replicates, with datasets merged where applicable, resulting in one dataset per sample.

#### HiFi Mapping

HiFi reads were mapped to the *Caenorhabditis elegans* reference genome^69^, using the pbmm2 tool based on minimap2. The PacBio data analysis was performed using SMRT Link version 13.0.0.207600. Parameters included: input minimum HiFi read quality of Q20, a minimum gap-compressed identity of 70%, and a minimum mapped length of 50 bp. The mean concordance of reads that mapped to the reference ranged from 98.51% to 99.32%. Approximately 338,158 CCS reads were mapped, with a 95th percentile read length of ∼15 kb and an average read length of 8.6 kb. The mean depth of coverage across the reference sequence was 26x.

#### Structural Variant Calling

HiFi reads (CCS BAM input) were aligned to the reference genome, and structural variation signatures were identified from the aligned BAMs. Variant calling and genotype assignment were conducted using the pbsv tool, detecting insertions, deletions, inversions, duplications, translocations, as both homozygous and heterozygous variants. Parameters included for variant calling were a min CCS Phred scale of 20, ≥10% read support in any one sample; ≥3 reads total; ≥500 bp).

#### Variant Calling

For each sample, mapped BAM files were merged using samtools (version 1.2.0) if multiple BAM files were available. Variants were called against the reference using DeepVariant (version 1.6.1) with default parameters and options “--model_type PACBIO --num_shards 12” (PMID: 30247488).

### Tc1 Transposon Mobilization Assay – Twitching Screen

To assess Tc1 transposon activity, a phenotype-based screen was performed. The insertion of Tc1 into the *unc-22* locus results in an observable twitching phenotype due to muscle spasms. To perform the assay a single nematode, confirmed to have no twitching phenotype, was isolated and propagated into a large population of approximately 300,000-500,000 individuals. The population was then synchronized using a bleach synchronization method.

Once the developmentally synchronized population reached the young adult stage, before egg-laying began, the worms were collected and washed with M9 buffer. Nicotine was added to a final concentration of 2%, which paralyzed all worms, allowing the twitching phenotype to become observable, as it is unaffected by nicotine.

The culture was screened under a dissecting microscope to identify twitching individuals. This allowed for the assessment of Tc1 transposon activity based on the presence of the twitching phenotype.

### Brood size quantification assay

Nematodes were developmentally synchronized by timed egg laying. Five gravid adult hermaphrodites were placed on individual S-plates for 3 hours and then removed. L4-stage progeny was selected and transferred individually to fresh S-plates for tracking. From the first day of adulthood, each worm was transferred daily to a new plate until reproduction ceased. Eggs laid were counted after each transfer, and hatched larvae and unhatched eggs were counted two days later. Brood size was calculated as the cumulative number of eggs laid across all reproductive days. Reproductive output was assessed as total eggs, viable progeny, and unhatched eggs. Quantifications were performed by visual inspection using a Leica M80 stereomicroscope. Statistical significance was calculated using two-tailed unpaired t-tests with Welch’s correction.

### RNA-seq processing

First, we preprocessed the raw samples with Fastp v0.20.0^70^ with the parameters -g -x -q 30 -e 30. The preprocessed samples were subsequently mapped with Salmon v1.1^71^ with the parameters –validateMappings –seqBias –gcBias and a decoy-aware reference genome based on WS281 including transposon transcripts from Wormbase^72^ .The mapped transcript-level counts were combined to the gene-level with tximport v1.14.2^73^ .filterByExpr was used to remove lowly expressed genes retaining 17674 genes for further analyses. edgeR v3.40.2^74^ was used for differential gene expression analysis. We identified three WT control outlier from batch 1, and 2 histamine treated WT outlier in a PCA based on sample clustering and removed them from subsequent analyses.

### Enrichment analyses

The 961 additively downregulated genes in the in heat-stressed WT, the AFD triple mutant, and heat-stressed AFD triple mutants were identified by selecting only those genes that were significantly (FDR<0.05) downregulated in all three pairwise comparisons. STRING v12^75^ was used for functional enrichment analysis, using all 17674 genes that remained after filtering as the statistical background. The histone modification enrichment analysis was done with the Enrichment Analysis from https://chip-atlas.org/^76^ for *C. elegans* (ce11), the experiment type “ChIP:Histone”, cell type class “Adult”, threshold for significance of 50, all significant up-regulated genes (FDR<0.05) as dataset A, all 17674 genes as dataset B, and a distance of −1kb <= TSS<= +1kb. Histone modifications for which more than 3 samples for whole worm samples that were over-enriched (fold change >1) exist were plotted. WormExp with a background gene set of all 17674 genes^77^ was used to compute enrichment for publicly available datasets. Gene set enrichment analysis (GSEA) for all transposable element transcripts, transposable element families, transcription factor motifs, and the sRNA-related genes were done with fgsea::fgseaMultilevel^78^. For the transcription factor motif analysis we downloaded all available curated motifs from the 10^th^ release (2024) of JASPAR^79^ and identified genes whose promoter region contained the motif with HOMER’s^80^ findMotifs.pl function and a motif score of at least 7. The identified genes were used for a GSEA.

### Public transcriptomic data analysis

The OP50-fed *inx-14 RNAi*^81^ data was downloaded from GSE216879. The processed ribozero emb-4(hc60) samples (GSE94455)^82^, and the lin-14 samples (GSE87523) were downloaded via Archs4^83^. The *atfs-1(et15)* data (GSE110984) was downloaded from the supplementary data of PMID30563508^48^. The hypoxia microarray dataset (GSE228851)^84^, and the *pmk-1(km25)* microarray dataset (GSE53732)^85^ were reanalyzed with GEO2R^86^. The pathogen PA14 dataset was downloaded from GSE122544^87^. The *hsf-1* samples (GSE241011) were downloaded from the supplementary data of PMID38895933^88^.

### WGS Filtering

We had to omit one of the F30 AFD triple mutant line that was WT for the gcy alleles and thus not a triple mutant.

To identify *de novo* F30 structural variants (SV) we filtered variants of each F30 generation sample that appeared in the respective P0 sample. For this we checked if the genomic region of each F30 SV is overlapping with any SV in the corresponding P0 sample. For imprecisely called SVs we extended the area of overlap by the length of the SV.

### Circos plots

The R v4.2.2 package circlize^89^ v.0.4.16 was used to draw the circos plots for all *de novo* F30 translocations (BNDs).

### Background mutation analysis

Genomic variants, including single nucleotide variants(SNVs), small insertions and deletions(INDELs), and structural variants (SVs), were annotated with SnpEFF 4.3t^90^ to assess the potential impact of each variant on gene function, allowing for the identification of possible background mutations.

### Transposable element insertions

The inserted sequences for all homozygous *de novo* F30 insertions were used as input for a BLASTN^91^ query against the known TE sequences from Wormbase (WS279.transposons.fa). TEs were only considered to match if the alignment length was bigger than 100. If for a query multiple TEs were matched we used the TE with the highest ‘bit score’.

### Repeat elements near break points

All *de novo* F30 break points (BNDs) were annotated to their nearest known repeat element in the ce11 genome. The known repeat elements were downloaded from the chromOut.tar.gz file from https://hgdownload.soe.ucsc.edu/goldenPath/ce11/bigZips/. Only repeats that were within 10bp of the break point were considered.

### CeNDR analysis

We downloaded the 20231213_c_elegans_transposon_calls.bed file from the *C. elegans* Natural Diversity Resource (CeNDR) https://caendr.org/data/data-release/c-elegans/latest ^92^. This file contains the count of several transposable element classes within 152 isotypes for *C. elegans*. For each isotype, the proportion of each transposable element family was calculate by dividing the number of TEs in each family by the total number of TEs identified in that strain.

### Data quantification, handling and statistical analysis

All data are presented as mean ± standard deviation (SD) or indicated in the figures and corresponding figure legends. The number of independent experiments (ℕ), biological replicates (N), and individual organisms (nematodes, n) are specified in the figures and corresponding figure legends. The specific statistical tests applied are detailed in the figure legends, with p-values reported directly in the diagrams. Boxplots are shown with the center line depicting the median value, the box limits showing the top and bottom quartiles, and the whiskers showing the 1.5x interquartile range. Linear regression plots are shown with the 95% confidence interval. Data handling, statistical testing, and plotting were conducted using Microsoft Excel 2019, R version 4.3.2, GraphPad Prism 9, and with Seaborn-0.11.0^93^ and Matplotlib-3.3.0^94^ in Python 3.6.10.

### *C. elegans* strains used for this study

| Strain | description | Genotype | origin |
| --- | --- | --- | --- |
| N2 Bristol | wild type |  | CGC |
| GN112 | AFD split caspase ablated - AFD:SCA | *pgIs2 [gcy-8p::TU#813 + gcy-8p::TU#814 + unc-122p::GFP + gcy-8p::mCherry + gcy-8p::GFP + ttx-3p::GFP]*. | CGC |
| BFF96 | AFD neuron specific transgenic histamine gated chloride channel expressing strain | *pgcy-8::HisCl1::SL2::mCherry::rab-3 3'UTR, pgcy-8::GCaMP6::rab-3 3'UTR; rol-6(GOF)* | Rechavi lab |
| BJS1062 | The integrated version of the BFF96 - AFD:HisCl1 | *pgcy-8::HisCl1::SL2::mCherry::rab-3 3'UTR, pgcy-8::GCaMP6::rab-3 3'UTR; rol-6(GOF)* | Generated by us |
| PY9248 | AFD triple mutant | *gcy-8(oy44); gcy-18(nj38); gcy-23(oy150) IV.* | Rechavi lab |
| BJS1456 | CRISPR regeneration of the PY9248 strain | *gcy-8(sbj126); gcy-18(sbj125); gcy-23(sbj127) IV* | Generated by us |
| IK597 |  | *gcy-8(oy44); gcy-18(nj38);gcy-23(nj37) IV* | CGC |

### Primers used for this study

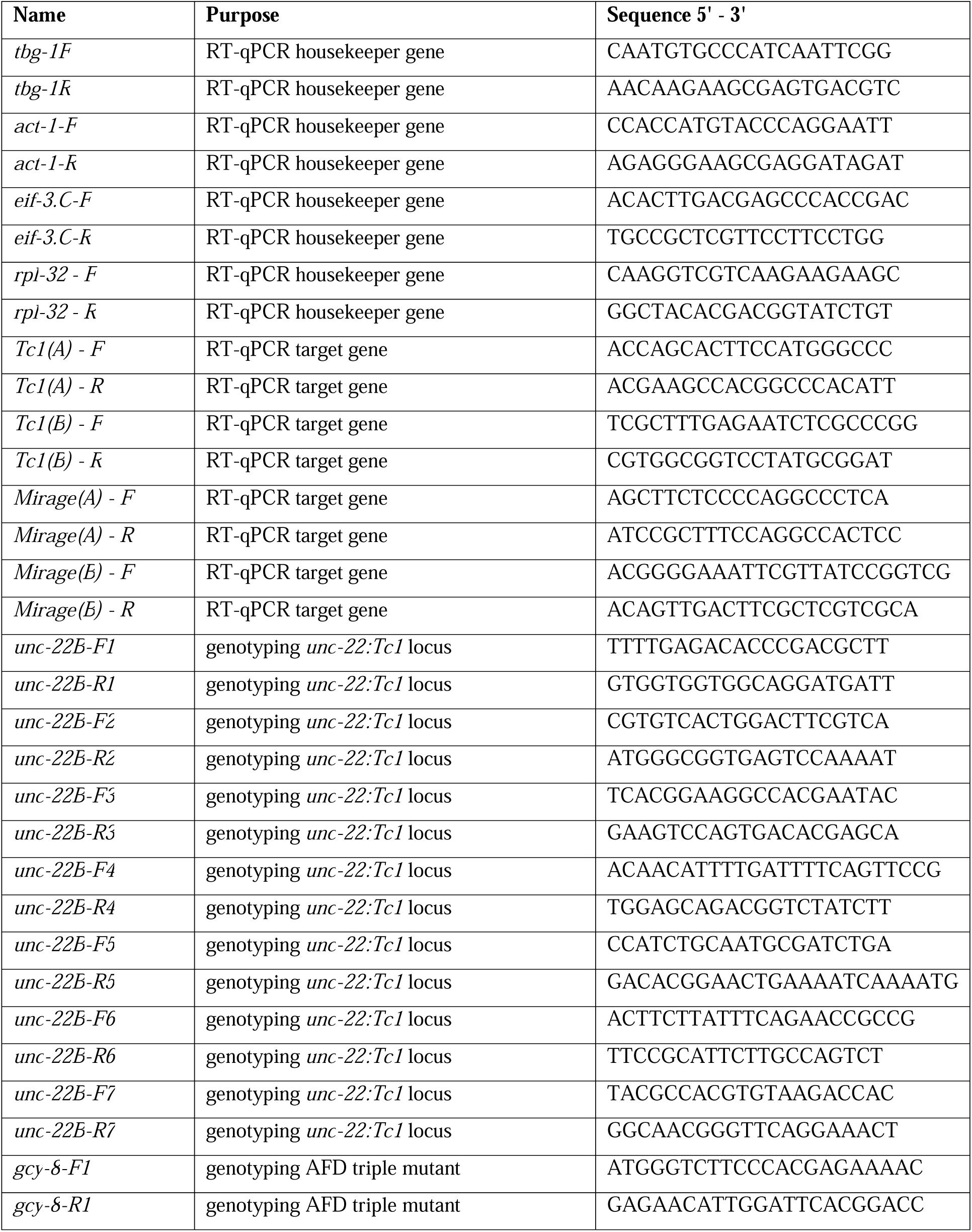
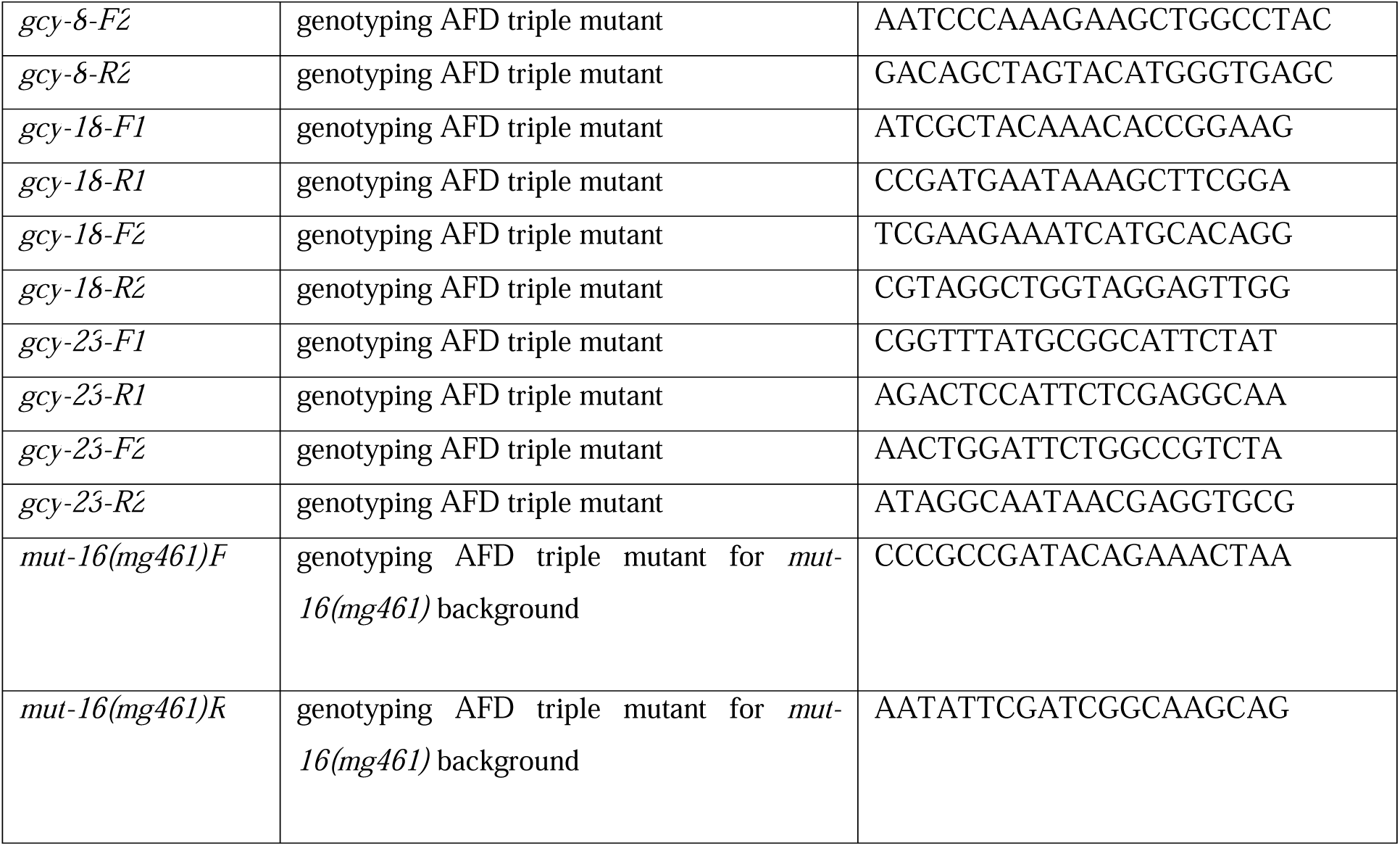

### Key resources, consumables, instruments used for this study

| Reagents: | Source | Identifier |
| --- | --- | --- |
| Nematode growth medium (NGM) agar | <sup>66</sup> |  |
| M9 buffer | <sup>66</sup> |  |
| Trizol® reagent | Invitrogen | 15596018 |
| Bromochloropropane (BCP) | Sigma-Aldrich | B9673 |
| SuperScript® III Reverse Transcriptase | Invitrogen | 18080085 |
| PowerUp™ SYBR™ Green Master Mix | Thermo Fisher Scientific | A25742 |
| Histamine dihydrochloride | Sigma | H7250-10G |
| Nicotine | Sigma | N3876 |
| Materials: |  |  |
| qPCR plates (96 x 0.1 ml plate) | BIOplastics | B60101 |
| 200 µl wide-bore pipette tips | Merck | AXYT205WBCRS |
| 1 ml wide-bore pipette tips | Merck | AXYT1005WBCRS |
| E. coli OP50 strain | CGC | N/A |
| E. coli NA22 strain | CGC | N/A |
| QIAGEN Puregene Cell and Tissue Kit | QIAGEN | 158388 |
| Qiagen RNeasy Mini Kit® | QIAGEN | 175032139 |
| **Instruments and Machines:** |  |  |
| Biobeam 8000 | Eckert & Ziegler,<br>Gamma Service Medical<br>GmbH |  |
| Centrifuge | Eppendorf | 5424 R |
| Nanodrop spectrophotometer | Thermo Scientific | N/A |
| CFX Connect real-time PCR detection system | BioRad | #1855201 |
| CFX96 Touch real-time PCR detection system | BioRad | #1854096 |
| Agilent TapeStation System | Agilent | Agilent |
| Dissecting microscope | Leica | CLS 150X |
| Axio Imager.M2 | Carl Zeiss | N/A |
| **Softwares:** |  |  |
| CFX Manager Software | BioRad | #1845000 |
| Microsoft Excel 2019 | Microsoft |  |
| R version 4.3.2 |  |  |
| GraphPad Prism 9 | GraphPad |  |

### Declaration of generative AI and AI-assisted technologies in the writing process

During the preparation of this work the authors used OpenAI ChatGPT-4o (2023) in order to grammatically proofread the text. After using this tool/service, the authors reviewed and edited the content as needed and take full responsibility for the content of the published article.

## Data availability statement

The RNA-seq data are available from the Gene Expression Omnibus (GEO) under the accession number GSE282988 and can be accessed with the reviewer password evcjgkgqdlmxdqz.

The proteomics data used in this study have been deposited to the PRIDE database with the accession number PXD056414.

The whole genome sequencing data are available in the BioProject database under the accession number PRJNA1190919 and can be accessed with https://dataview.ncbi.nlm.nih.gov/object/PRJNA1190919?reviewer=14ujg5s97i19vhb8u0plfre30 8.

Each figure representing an analysis has corresponding raw data and statistical comparisons; see the statistical source data supplementary file.

**Supplementary Figures Supplementary Figure 1.**

A) RT-qPCR primer binding sites for transposase mRNA measurements. Two independent primer pairs targeting the same transposase consensus sequence in their respective TE families were used.

A For the Tc1 transposase, the green primer pair Tc1(A) and the blue primer pair Tc1(B) were used. For the Mirage transposase, the yellow primer pair Mirage(A) and the red primer pair Mirage(B) were employed.

RT-qPCR was performed to quantify the differential expression of transposase mRNA in various AFD neuron-depleted *C. elegans* strains. The experiment was repeated three times, each time with independently set up three independent cohorts consisting of 3000 nematodes. All biological replicate was measured in technical triplicates. Each data point represents the mean of a biological replicate, with data from these nine independent populations combined (=3, N=9). Target transposases, were measured with: Tc1(A) for Tc1 transposase, and Mirage(A) for Mirage transposase. Relative quantities were calculated using the 2ΔCt method and normalized using a normal factor derived from the relative quantities of the housekeeping genes *tbg-1*, *act-1*, *eif-3.C*, and *rpl-32*. This provided the normalized fold change (FC) values for the target transposases. Statistical significance was determined using an unpaired t-test with Welch’s correction, following normality assessment with the Shapiro-Wilk test. The p-values for the comparisons are indicated on the figure, along with the percentage change where relevant.

B) Differential expression of transposase mRNAs in AFD split caspase-depleted strain (AFD:SCA) and wild type in comparison.

C) Differential expression of transposase mRNAs in AFD histamine-gated chloride channel strain (AFD:HisCl1) and wild type in comparison. Populations were cultured on standard NGM and NGM supplemented with 1 mM histamine.

D) Differential expression of transposase mRNAs in AFD triple mutant strain compared to the wild type control, cultured under normal experimental temperatures and heat-stress conditions.

**Supplementary Figure 2.**

Genotyping of individual worms using single-worm PCR for the AFD triple mutant in the *mut-16(mg461*) background, compared to a wild-type control. PCR was performed on three randomly selected nematodes (N=3) from their respective population. The expected PCR product size for the wild-type allele is 824 bp, while the expected product size for the *mut-16(mg461)* allele is 373 bp.

**Supplementary Figure 3.**

A) Overview of gene set enrichment analyses (GSEA) for transposable elements in the annotated pairwise RNA-seq comparisons. The y-axis shows the normalized enrichment score of a GSEA. A positive value indicates the enrichment of transposable element genes in the upregulated genes of the respective pairwise comparison. The underlying results are shown in Supplementary Figure 3B-G *<0.05, **<0.01, ***<0.001, ns>=0.05.

B) GSEA of the transposable element genes in the WT heat stress vs. control RNA-seq comparison. The y-axis shows the enrichment score. The x-axis the gene rank, with the highest expressed gene being rank 0. TE genes are shown as lines on the bottom. The red dashed lines show the peaks of the running sum of the enrichment score (green line).

C) The same as B) but for the comparison AFD triple mutant vs WT.

D) The same as B) but for the comparison AFD triple mutant heat stress vs AFD triple mutant control.

E) The same as B) but for the comparison AFD triple mutant heat stress vs WT heat stress.

F) The same as B) but for the comparison AFD:SCA vs WT

G) The same as B) but for the comparison AFD:HisCl1 histamine vs WT control.

H) Heatmap of the average logFC of the transposable element classes of the respective RNA-seq comparisons. The color code shows the average logFC.

I) Heatmap of the normalized enrichment score of the transposable element classes with at least 5 genes. The color code shows the normalized enrichment score calculated via a GSEA. The hierarchical clustering of the columns was done with a correlation metric and the Ward method.

**Supplementary Figure 4**

A) The distribution of genes significantly downregulated (blue) or upregulated (red) in the constitutive active *crh-1(tz2)* mutant in the differentially expressed genes in the comparison AFD triple mutant vs. WT. The x-axis shows the logFC of the AFD triple mutant. The density plots show the distribution of the genes that are significantly upregulated (red), respective downregulated (blue), in the *crh-1(tz2)* dataset.

B) The same as A) but for the *pmk-1(km25)* dataset.

C) The same as A) but for the *emb-4(hc60)* dataset.

D) The same as A) but for the *miR-243(n4759)* dataset.

E) The same as A) but for the *atfs-1(et15)* dataset.

**Supplementary Figure 5**

Heatmap of the public datasets shown in Figure 3F. All genes from the sRNA regulation pathway (Figure 2D) and genes known to be affected by the AFD neuron (Figure 3A) are shown. The color code shows the logFC. The hierarchical clustering was done with a correlation metric and the Ward method.

**Supplementary Figure 6**

Single GSEA plots of the sRNA regulation related genes for Figure 3F.

A) AFD triple mutant vs. WT RNA-seq comparison. The y-axis shows the enrichment score. The x-axis the gene rank, with the highest expressed gene being rank 0. sRNA regulation related genes are shown as lines on the bottom. The red dashed lines show the peaks of the running sum of the enrichment score (green line).

B) The same as A) but for the public dataset *lin-13(n770)* vs WT.

C) The same as A) but for the public dataset *flp-6(ok3056)* vs WT.

D) The same as A) but for the public dataset *inx-14* RNAi vs Control.

E) The same as A) but for the public dataset *emb-4(hc60)* vs WT.

F) The same as A) but for the public dataset *atfs-1(et15)* vs WT.

G) The same as A) but for the public dataset Pseudomonas aeruginosa (PA14) vs Control.

H) The same as A) but for the public dataset Hypoxia vs Control.

I) The same as A) but for the public dataset *pmk-1(km25)* vs WT.

J) The same as A) but for the public dataset *hsf-1(sy441)* vs WT.

**Supplementary Figure S7.**

Brood size of AFD neuron-depleted strains:AFD triple mutant, AFD:SCA A), and AFD:HisCl1 with and without 1 mM histamine treatment B), compared to their corresponding wild-type controls. Each data point represents the total progeny of a single nematode throughout its reproductive lifespan. All recordings were performed simultaneously and are shown in separate panels (A and B); note that the non-treated wild-type values are shared across both panels. Statistical significance was calculated using a two-tailed unpaired t-test with Welch’s correction. P-values for comparisons against the non-treated wild type are indicated above each dataset, while p-values for the comparison between histamine-treated wild type and AFD:HisCl1 are shown above the connecting lines.

**Supplementary Tables**

**Supplementary Table 1:** STRING-DB analysis of the additively downregulated genes

**Supplementary Table 2:** WormExp Analysis of the signficantly upregulated genes in the comparison AFD triple mutant vs WT

**Supplementary Table 3:** SVs impacting transposon related genes

**Supplementary Table 4:** SNVs and small INDELs impacting transposon related genes. Supplementary **Table 5**: Analysis of active transposable element families in isotypes of C. elegans

