## Supplementary figures and images for "Thermosensory neuron dysfunction alters transposable element regulation"

### Supplementary Figure 1

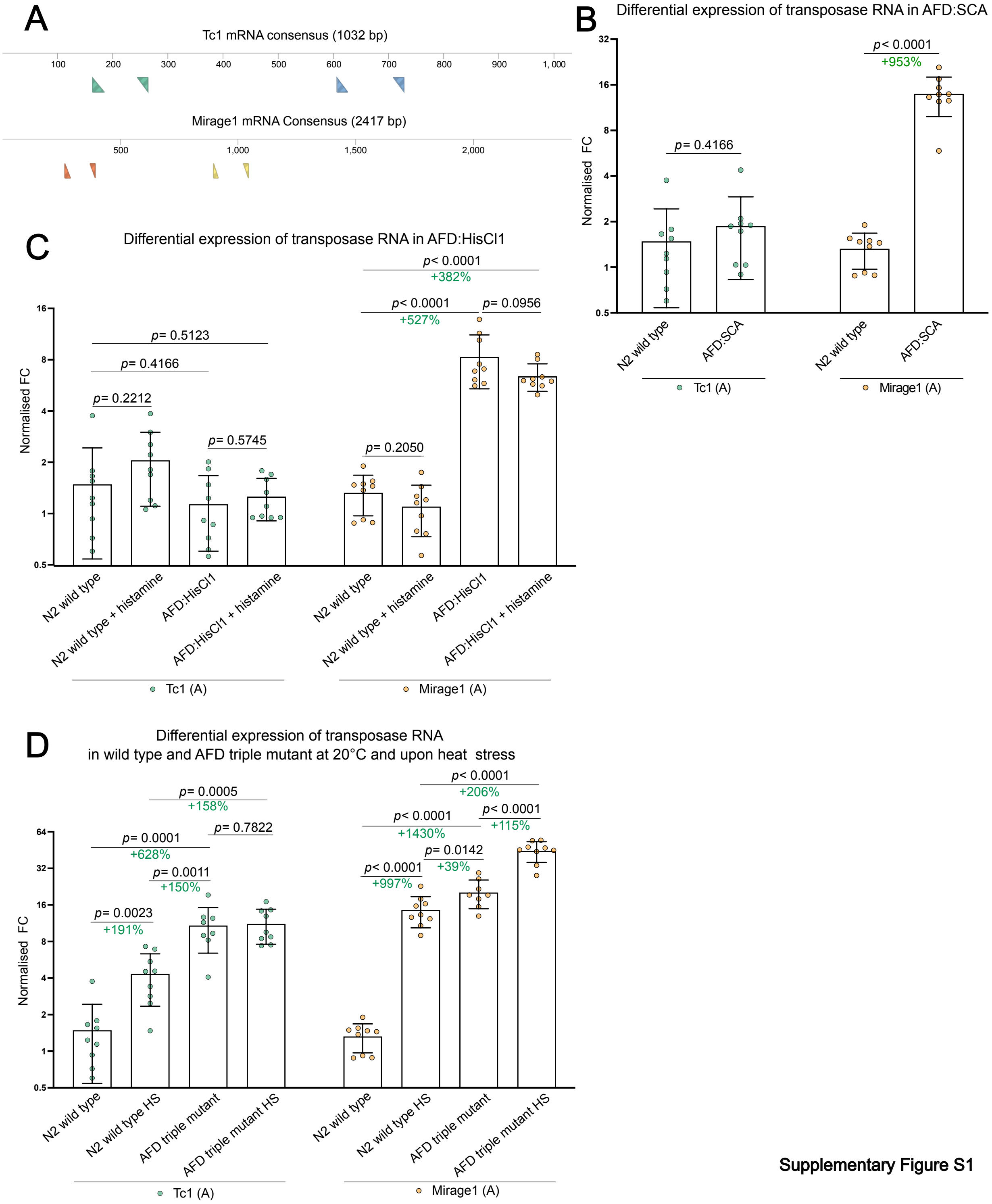


**Supplementary Figure S1**

### Supplementary Figure 2

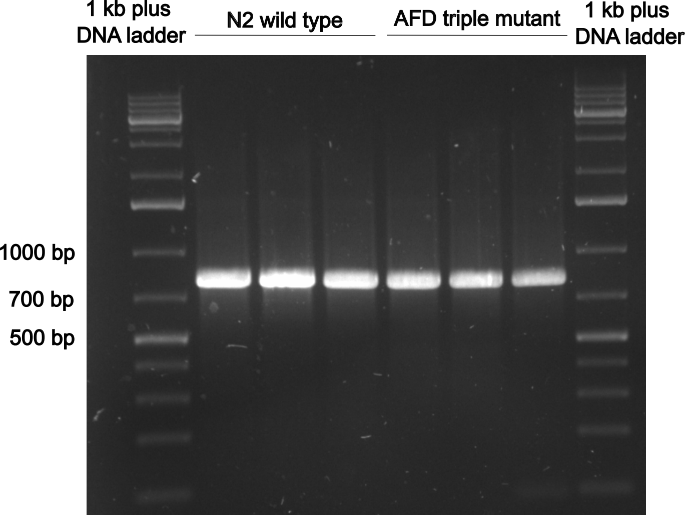


Supplementary Figure S2

### Supplementary Figure 3

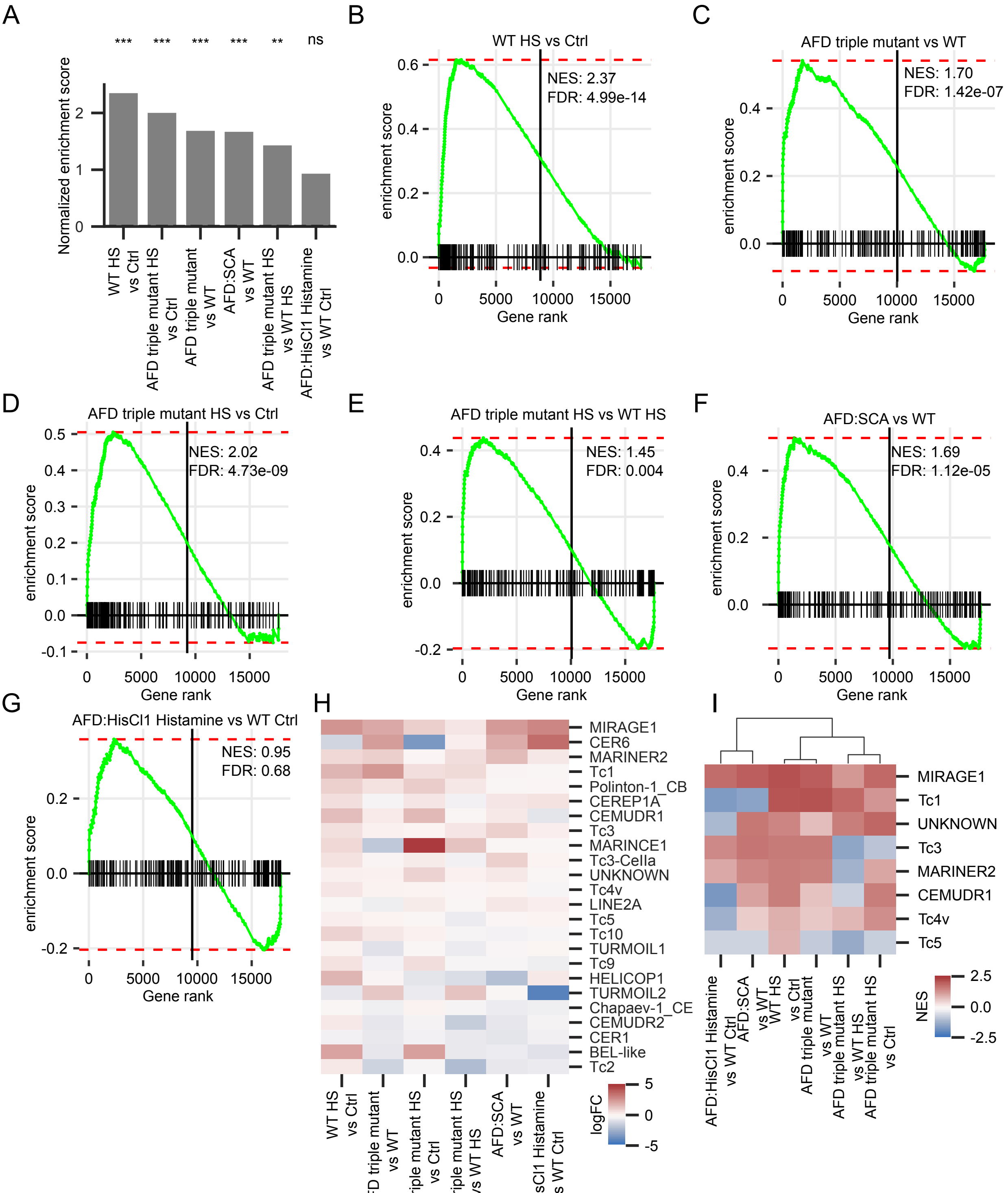


**Supplementary Figure S3**

### Supplementary Figure 4

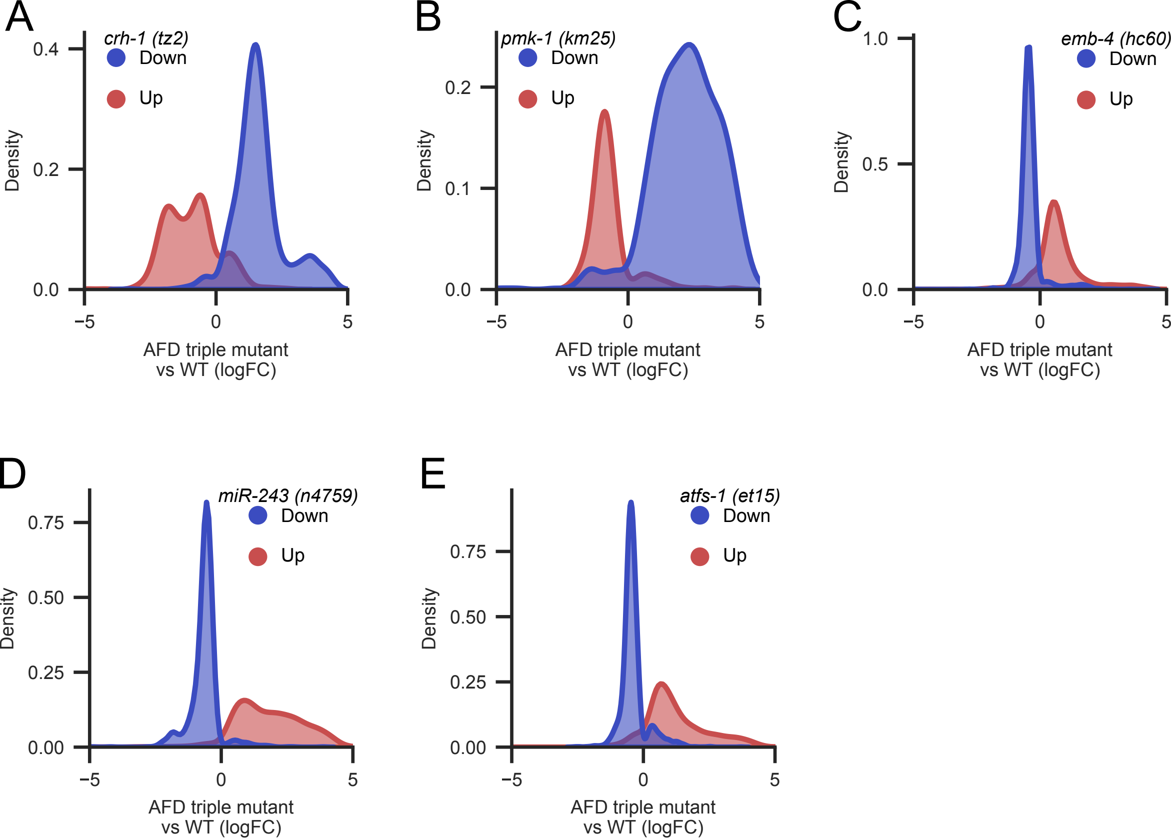


**Supplementary Figure S4**

### Supplementary Figure 5

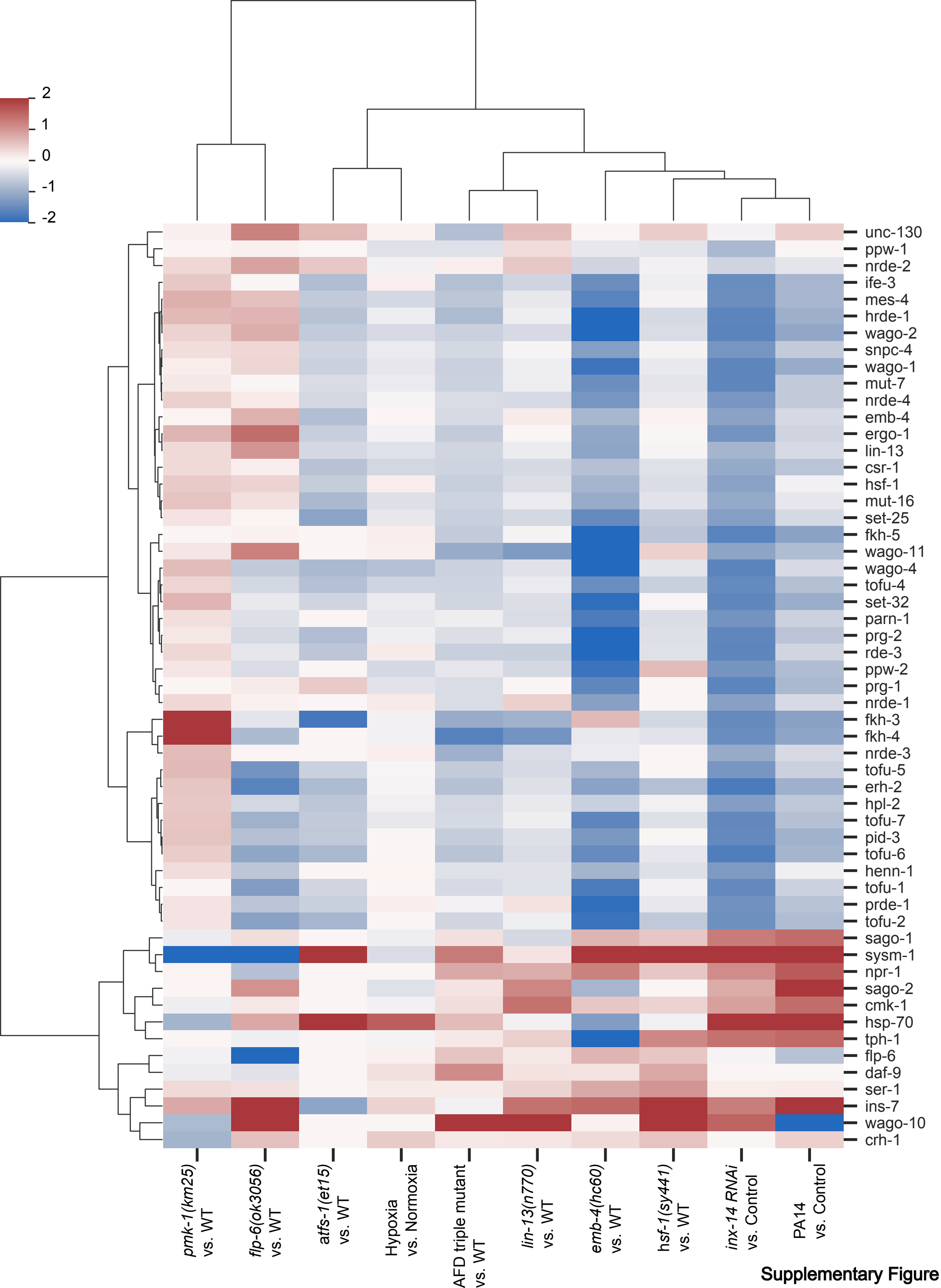


**Supplementary Figure S5**

### Supplementary Figure 6

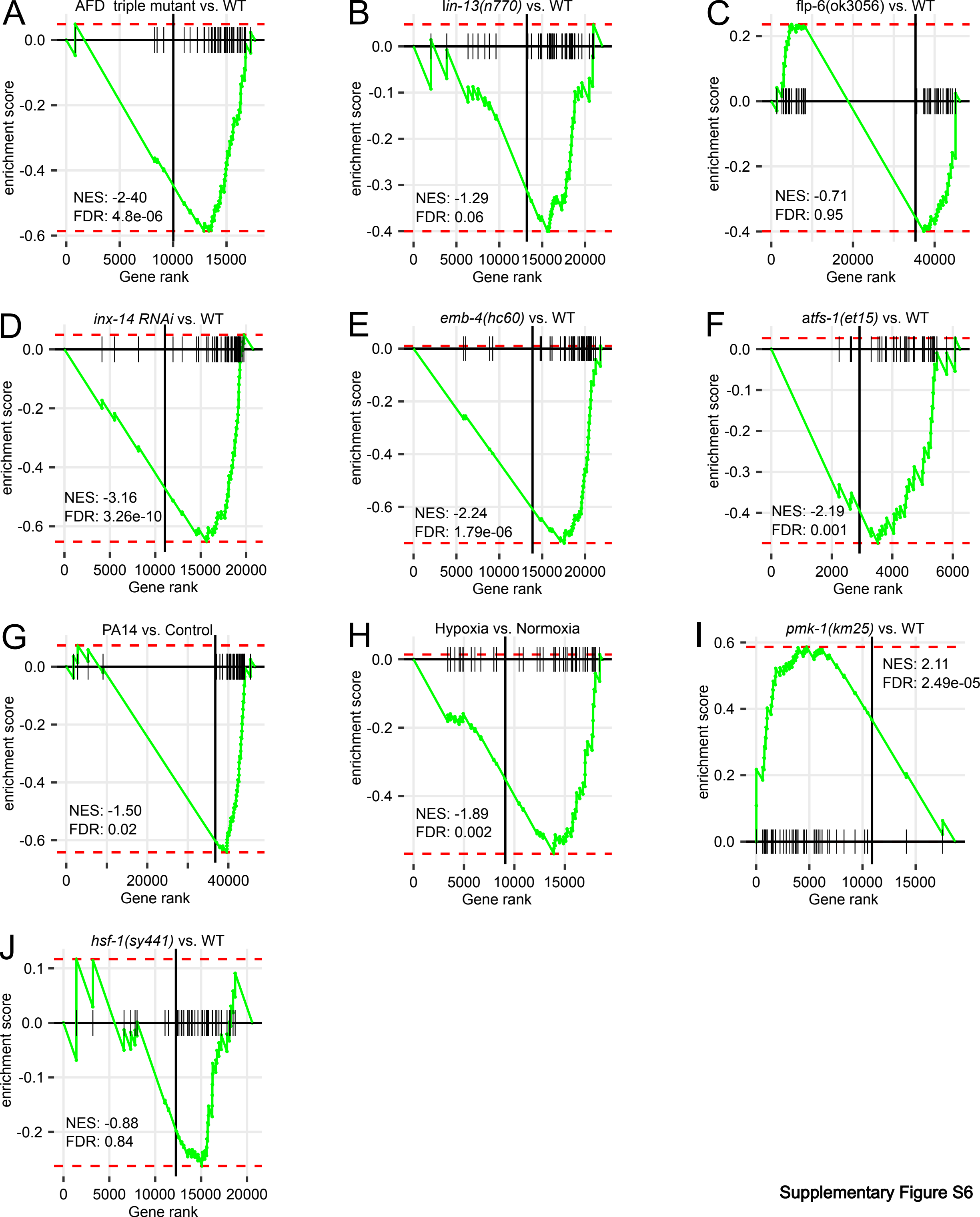


Supplementary Figure S6

### Supplementary Figure 7

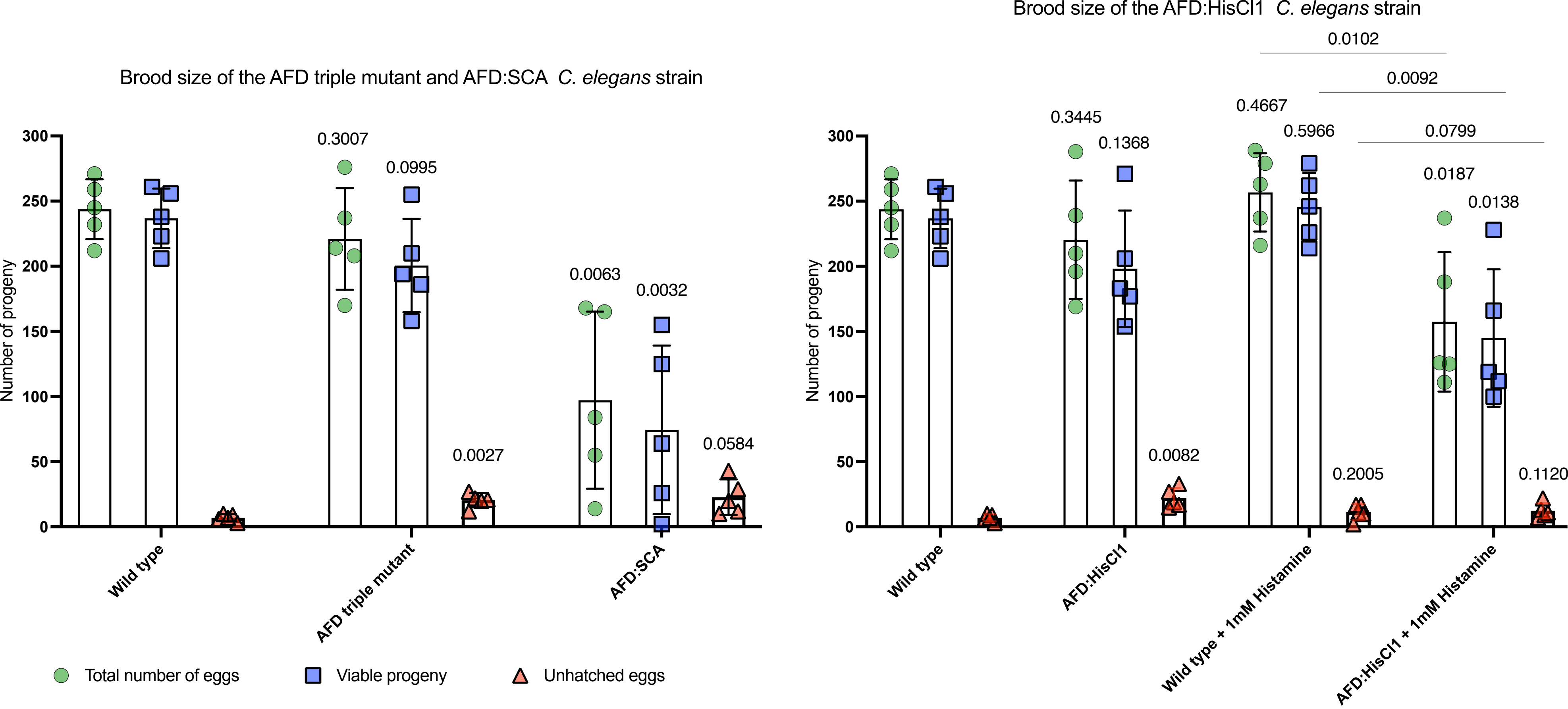


Supplementary Figure S7
